# Coevolution of neoplastic and non-neoplastic reactive astrocyte states converges on mesenchymal-like and injury-response programs during murine glioblastoma progression and post-radiotherapy recurrence

**DOI:** 10.64898/2025.12.18.695184

**Authors:** David Lindgren, Rebecca Rosberg, Karolina I. Smolag, Elinn Johansson, Dimitra Manou, Jonas Sjölund, Sebastian Braun, Bengt Phung, Katja Harbst, Pauline Jeannot, Maria Malmberg, Crister Ceberg, Göran B. Jönsson, Kristian Pietras, Alexander Pietras

**Author notes:** These authors contributed equally (David Lindgren, Rebecca Rosberg). Corresponding author: Alexander Pietras.

## Abstract

Glioblastomas initially respond to radiotherapy but invariably recur, often within high-dose radiation treatment fields. Although stromal radiation responses are incompletely understood, evidence suggests that the tumor microenvironment becomes tumor-supportive after therapy. Using a genetically engineered glioblastoma mouse model, we profiled healthy brain, primary tumors, and post-radiotherapy recurrences with single-cell and spatial transcriptomics and immunohistochemistry. Across 13 non-neoplastic cell types and 10 tumor cell states, we mapped transcriptional adaptations accompanying progression from healthy brain to primary and recurrent glioblastoma. We identified distinct astrocyte states linked to disease stage, including reactive non-neoplastic astrocytes and tumor cells adopting reactive astrocyte-like phenotypes. Reactive astrocyte-like tumor cells with mesenchymal and injury-response signatures were enriched after radiotherapy and persisted in recurrent tumors. Receptor–ligand interactions between reactive astrocytes and tumor cells included known and putative drivers of aggressiveness. These findings highlight convergent reactive astrocyte programs in astrocytes and tumor cells as potential mediators of glioblastoma radioresistance.

## Introduction

Glioblastoma (GBM) remains one of the most challenging cancers to treat, with median survival times of 12-15 months post-diagnosis and virtually no long-term survivors^1^. As GBMs progress, tumor cells co-evolve within an ever-changing microenvironmental landscape, and these tumors are characterized by remarkable heterogeneity both at the tumor cell level and at the level of various stromal and immune cell types. Recent efforts have established roles for the microenvironment in promoting aggressive cell states of highly plastic GBM cells; for instance, mesenchymal-like (MES-like) phenotypes of GBM cells are driven by interactions between tumor and immune cells^2,3^, cells displaying high stemness can be influenced by oxygen and nutrient availability in distinct tumor niches^4,5^, and increasing focus is directed towards the role of neuronal and neuroglial networks in changing GBM cell fates^6–8^.

Essentially all GBMs are treated with radiotherapy (RT), but tumors typically recur as difficult-to-treat lesions, frequently in brain areas overlapping with the areas exposed to high-dose irradiation during treatment of the primary tumor^9–11^. A detailed understanding of changes to the microenvironment during RT and progression to recurrent disease is lacking, but mounting evidence suggests that RT-induced adaptations of immune and stromal cells may generate tumor-supportive conditions in the post-RT brain microenvironment^12^. Specifically, tumor-associated astrocytes respond to RT with a reactive phenotype that has been associated with traits of aggressive tumor cells such as therapeutic resistance and increased invasive capacities^13^. Furthermore, astrocytes are immune-modulatory cells that signal reciprocally with macrophages and microglia, and thereby contribute to shaping the notoriously immune-suppressive microenvironment of GBM^14,15^. Astrocyte reactivity has emerged as a promising therapeutic target not only in brain tumors, but also in other neurological conditions^16^. While recent efforts have contributed to an increased understanding of the process of astrocyte reactivity in health and disease^17^, relatively little is still known about distinct astrocyte states present in GBM pre- and post-RT, and how these states relate to astrocytes in the healthy brain.

Here, we used a genetically engineered mouse model of GBM and sampled tissue from healthy brain, primary GBM, and post-RT recurrent GBM for single cell and spatially resolved transcriptomic analyses supplemented with immunohistochemical verifications. We detailed transcriptomic adaptations of at least 13 distinct non-neoplastic cell types and 10 tumor cell states, during progression from healthy murine brain to primary GBM and post-RT recurrent GBM. We found distinct astrocyte states associated with disease progression, some of which were non-neoplastic reactive astrocytes, and others that were dominated by tumor cells adapting a reactive astrocyte state. Tumor cells with a reactive astrocyte phenotype increased drastically within the surviving tumor cell population immediately after RT, and were enriched in post-RT recurrent GBM. Receptor-ligand complexes between reactive astrocytes and tumor cells enriched in recurrent tumors included known and novel drivers of aggressive GBM growth. Together, our findings establish phenotypic and functional heterogeneity among tumor-associated astrocytes in GBM, and underscore reactive astrocyte states as potential drivers of GBM radioresistance.

## Results

### A murine model of GBM initiation and post-radiotherapy relapse

To investigate how radiotherapy influences the glioblastoma (GBM) microenvironment, we generated murine GBMs using the RCAS/tv-a system to deliver human PDGFB (hPDGFB), a short hairpin RNA targeting Tp53 (shTrp53), and red fluorescent protein (RFP) to Nestin-expressing cells in the neonatal mouse brain, as previously described^18,19^. Samples were collected from healthy brain tissue, resected primary (non-irradiated) tumors, and post-radiotherapy recurrent tumors that developed in symptomatic mice following cranial irradiation with 10 Gy (Figure 1A). Radiotherapy extended the median survival of tumor-bearing mice from 32.5 days to 55 days (Figure 1B). Immunofluorescence analysis revealed RFP⁺/Olig2⁺ tumor cells intermingled with Gfap⁺ astrocytes in GBM samples, whereas no RFP⁺ cells were detected in healthy brain tissue (Figure 1C).

**Figure 1.**
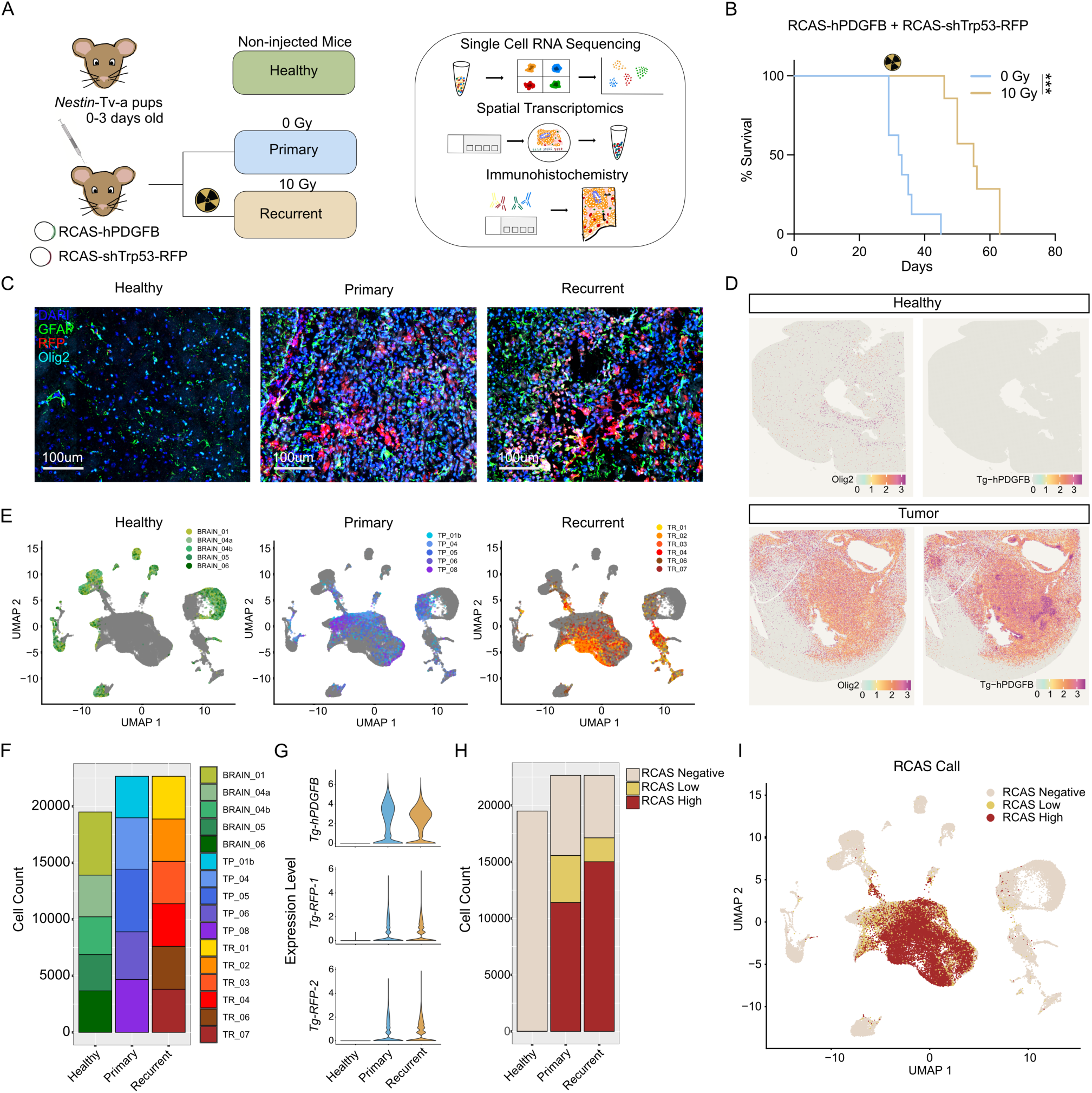
A murine model of GBM and post-radiotherapy recurrence. **(A)** Schematic of the study design and analyses. Newborn Nestin-tv-a mice received RCAS-hPDGFB and RCAS-shTrp53-RFP viral injections. Tissue cell biopsies were collected from primary tumor and post-radiotherapy recurrent stages as well as from healthy controls. Tissue samples were processed for scRNA-seq, Visium spatial transcriptomics, and immunohistochemistry. **(B)** Kaplan–Meier survival curves for tumor-bearing mice treated with 0 Gy or 10 Gy cranial irradiation. Radiotherapy significantly prolonged survival. **(C)** Immunofluorescence images of healthy, primary, and recurrent brain tissue showing DAPI, GFAP, RFP, and Olig2. RFP⁺ tumor cells are abundant in primary and recurrent tumors but absent in healthy brain. **(D)** Spatial expression maps (Visium HD) showing Olig2 and Tg-hPDGFB expression in healthy and tumor sections. **(E)** UMAP projections of 64,804 scRNA-seq profiles from 5 healthy, 5 primary, and 6 recurrent samples. Non-neoplastic cell types form multiple clusters, some enriched for specific sample types. **(F)** Sample contributions to the scRNA-seq dataset across healthy (BRAIN), primary (TP), and recurrent (TR) groups. **(G)** Violin plots showing Tg-hPDGFB and Tg-RFP expression restricted to cells from tumor tissues. **(H)** Proportions of RCAS^neg^, RCAS^low^, and RCAS^high^ cells across healthy, primary, and recurrent samples. **(I)** UMAP projection of cells labeled with the RCAS-based classification.

Whole-transcriptome profiling was performed on tumor and healthy brain tissues using single-cell RNA sequencing (scRNA-seq; 10x Genomics Flex) and spatial transcriptomics (10x Genomics Visium v1 and Visium HD). Spatial transcriptomic analysis confirmed the co-expression of *hPDGFB*, *RFP*, and *Olig2* in tumor tissue (Figure 1D and Supplementary Figure 1A). Dimensionality reduction using UMAP of 64,804 cells from the scRNA-seq dataset separated cells into multiple, sample-independent clusters, many of which contained cells from all sample types, while some were composed almost exclusively of cells from healthy brain tissue or dominated by a single tumor type (Figure 1E-F). Expression of *hPDGFB* and *RFP* was exclusive to tumor samples and cells expressing these transgenes formed a large, distinct cluster (Figure 1G and Supplementary Figure 1B). To aid identification of neoplastic cells, we constructed an RCAS-signature based on expression of the transgenes and cells were classified as RCAS^high^, RCAS^low^, or RCAS^neg^. Within tumor samples, 65% and 69% of cells in primary and recurrent tumors, respectively, were classified as RCAS^high/low^ (Figure 1H-I, Supplementary Figure 1C-G).

### Single-cell transcriptomic characterization of mouse GBM and tumor microenvironment

We next performed graph-based SNN clustering of the scRNA-seq dataset, resulting in 26 transcriptionally distinct clusters across the 64,804 cell transcriptomes (Figure 2A). Integration of cluster identity with RCAS-transgene expression showed that these clusters segregated into three major compartments (Figure 2B): (i) a large group containing exclusively RCAS^neg^ cells from all sample types, (ii) a mixed group containing both RCAS^neg^ and RCAS^low/high^ cells, and (iii) a group composed almost entirely of RCAS^low/high^ cells originating from tumor samples (Figure 2B and Supplementary Figure 2A-D). Based on this structure, we classified 30,345 cells as non-neoplastic, 33,727 as neoplastic, and 732 cells with ambiguous profiles (Figure 2C).

**Figure 2.**
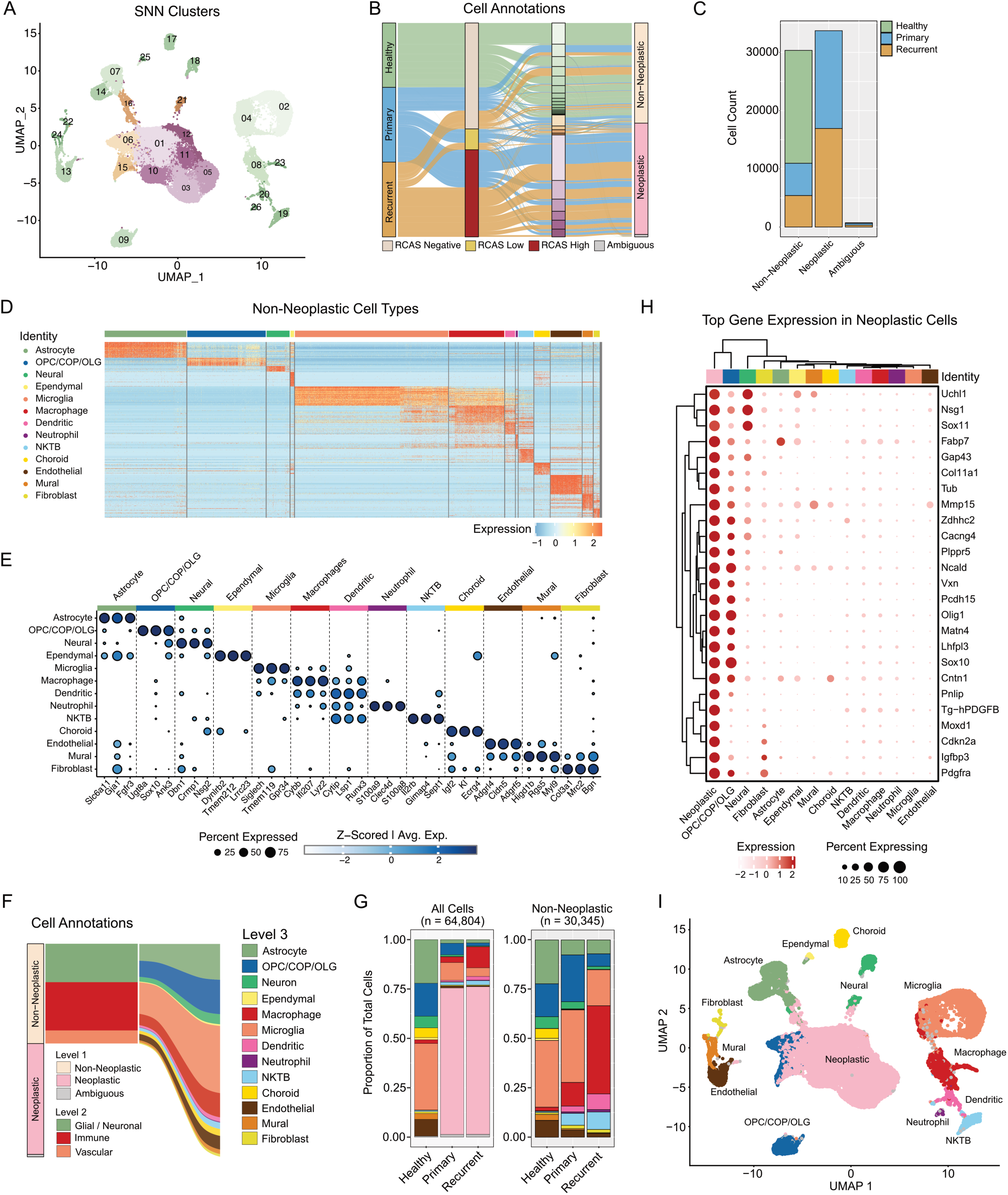
Single cell transcriptomic characterization of mouse GBM and tumor microenvironment. **(A)** UMAP projection of 64,804 scRNA seq profiles, showing 26 transcriptionally distinct clusters identified by graph based SNN clustering. **(B and C)** Sankey diagram linking sample type (Healthy, Primary, Recurrent) to RCAS transgene status (RCAS^neg^, RCAS^low^, RCAS^high^) and to the 26 SNN clusters, which are grouped into three broad compartments based on RCAS expression patterns (see Supplementary Figure 2), and final Level 1 classification. The flows illustrate how clusters segregate into non-neoplastic, neoplastic, or ambiguous compartments. Barplot with cell numbers and for each category and labeled according to sample type (C). **(D)** Heatmap showing marker gene expression across the 30,345 non-neoplastic cells after recursive re-clustering. Columns represent individual cells grouped by their assigned identity, and rows represent the top 50 marker genes for each major cell class. **(E)** Dot plot showing the top three defining marker genes for each major cell population. Dot size reflects the proportion of cells expressing each gene within the indicated group, and color encodes the average scaled expression (Seurat DotPlot metrics). **(F)** Sankey diagram showing the flow from Level 1 compartments (non-neoplastic, neoplastic, ambiguous) to Level 2 classes (glial/neural, immune, vascular), and their further resolution into Level 3 cell type identities defined in Figure 2D–E. **(G)** Relative abundance of major cell types among all cells (left) and within non-neoplastic cells (right). Recurrent tumors exhibit expansion of infiltrating myeloid populations and reduced microglia. **(H)** Clustered dot plot displaying the top 25 genes that distinguish neoplastic from non-neoplastic compartments. Dot size indicates the percentage of cells expressing each gene, and dot color reflects scaled expression values. Rows (genes) and columns (cell groups) are hierarchically clustered. **(I)** UMAP projection of all 64,804 cells, using the same embedding as in Figures 1 and 2A but annotated with the final curated cell type assignments, providing an overview of the transcriptional landscape of the PDGFB driven mouse GBM model.

To resolve the cellular diversity of the tumor microenvironment, we re-clustered the 30,345 non-neoplastic cells in a recursive approach and annotated them using the Allen MapMyCells tool (RRID:SCR_024672). This analysis identified all major resident brain and stromal cell populations, including astrocytes (*Slc6a11*, *Gja1*), oligodendrocyte lineage cells (*Sox10*, *Ugt8a*), neuronal lineages (*Nsg2*, *Dynlrb2*), microglia (*Siglech*, *Tmem119*), bone-marrow-derived macrophages (*Cybb*, *Lyz2*), dendritic cells (*Runx3*), NK/T/B lymphocytes (*Il2rb*, *Gimap4*), neutrophils (*S100a8*), endothelial cells (*Cldn5*), mural cells (*Myl9*), fibroblasts (*Col3a1*), and choroid plexus epithelium (Figure 2D-2E). The contribution of major compartments across all sample types is summarized in Figure 2F.

Although these identities were represented across all conditions, their proportions changed markedly between healthy tissue, primary tumors, and post-radiotherapy recurrent tumors (Figure 2G). Immune populations such as bone-marrow-derived macrophages, dendritic cells, neutrophils, and NK/T/B cells increased substantially in tumor samples, with the most dramatic expansion observed in recurrent tumors. In contrast, microglia, the resident CNS macrophages, decreased in relative abundance from 37% of non-neoplastic cells in primary tumors to 18% in recurrent tumors, consistent with progressive shift from microglia-dominant to macrophage-dominant myeloid populations (Figure 2G, and Supplementary Figure 2E). Even though individual non-neoplastic cell types varied between treated and untreated tumors, the relative abundance of neoplastic cells was nearly identical between treated and untreated tumors (74 percent versus 75 percent, Figure 2G).

With the major cell populations defined, we next examined how neoplastic cells diverge transcriptionally from their non-neoplastic counterparts. This comparison uncovered a set of tumor-specific programs enriched in the neoplastic compartment (Figure 2H), including increased expression of lineage plasticity regulators (*Sox11*, *Fabp7*), oligodendrocyte lineage genes (*Olig1*, *Sox10*), PDGF pathway components (*Pdgfra*), ECM remodeling factors (*Col11a1*, *Mmp15*), and markers of metabolic adaptation (*Igfbp3*, *Moxd1*).

In summary, single-cell RNA-seq delineated the diverse cellular composition of the PDGFB-driven mouse GBM microenvironment, identifying both tumor-intrinsic and microenvironmental changes associated with disease progression and recurrence. A final UMAP projection, now annotated with the refined cell type identities, provides an overview of the transcriptional landscape of all 64,804 cells (Figure 2I).

### Composition of astrocytes in the tumor microenvironment

Astrocytes are increasingly recognized as important regulators of the GBM microenvironment, particularly after irradiation where reactive astrocyte states have been linked to tumor aggressiveness and recurrence ^13,20–22^. To characterize these populations, we performed focused analyses on the 5,110 astrocytes identified from healthy and non-neoplastic tumor compartments. Graph-based clustering segregated the astrocytes into three transcriptionally distinct subclusters, two of which were dominated by cells from healthy tissues (Figure 3A–B). All astrocyte subclusters expressed canonical pan-astrocyte markers such as *Slc6a11*, *Gja1*, *Aqp4*, and *Aldoc*^23^, with spatial expression patterns matching those of astrocytes, thus confirming the cellular identity of this cell cluster (Figure 3C–D). Annotation of the two healthy-tissue–dominated clusters using the Allen MapMyCells^24^ tool classified them as likely telencephalic (TE) and non-telencephalic (NT) astrocytes, respectively, indicating that these subclusters reflect regional specialization of astrocytes across anatomical territories. This was corroborated by Visium HD spatial mapping, which revealed distinct regional expression patterns for module scores derived from TE- and NT-enriched gene sets (Figure 3C).

**Figure 3.**
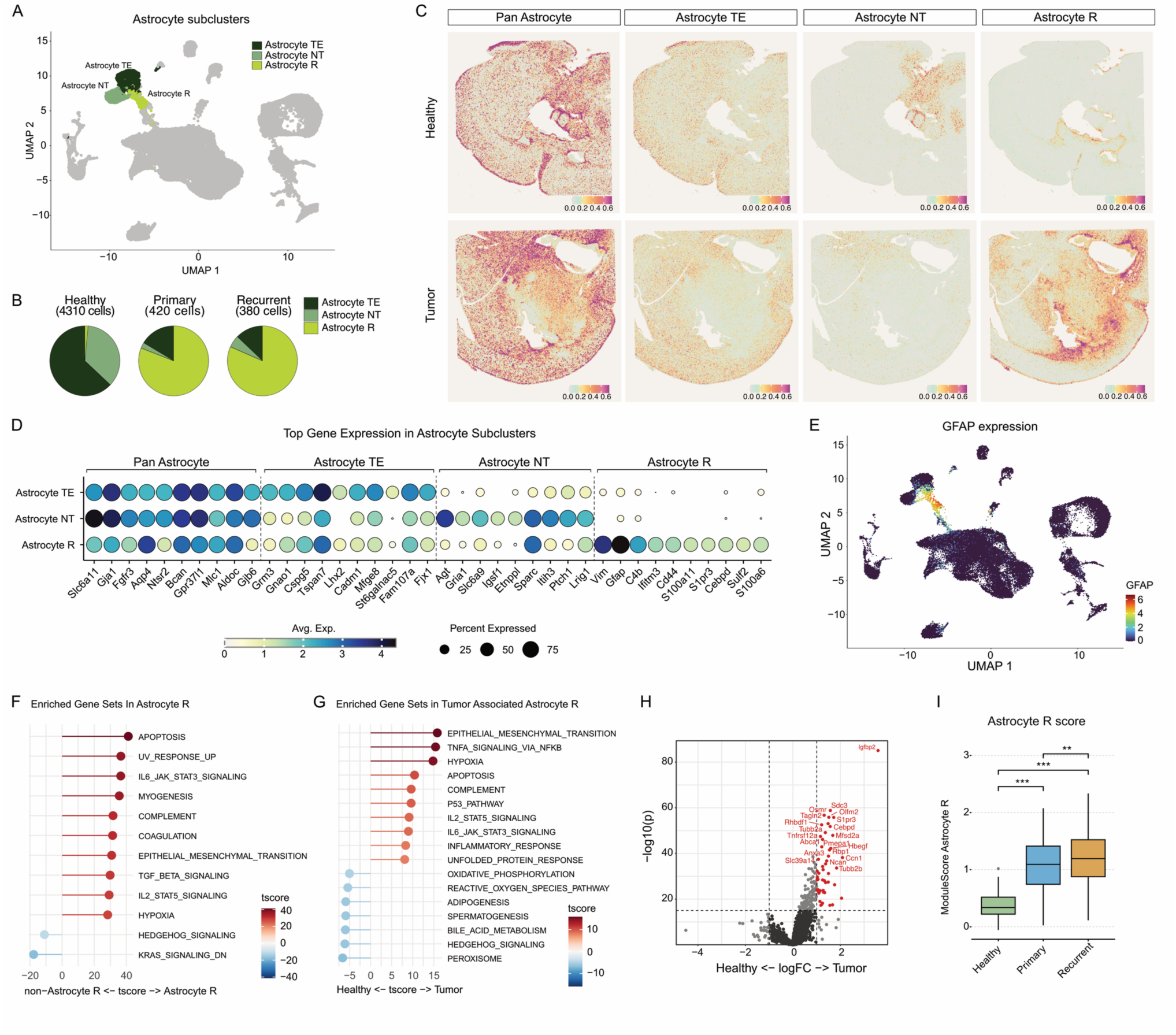
Astrocyte lineage diversification and reactive-state induction in murine GBM. **(A**) UMAP projection highlighting the astrocyte lineage after subsetting and re-clustering of 5,110 astrocytes. Three transcriptionally distinct astrocyte subclusters were identified: telencephalic astrocytes (Astrocyte TE), non-telencephalic astrocytes (Astrocyte NT), and reactive tumor-associated astrocytes (Astrocyte R). **(B)** Distribution of astrocyte subclusters across sample types. Healthy brain tissue is dominated by TE and NT astrocytes, whereas primary and recurrent tumors are highly enriched for the reactive astrocyte population (Astrocyte R). **(C)** Visium HD spatial mapping of astrocyte module scores derived from scRNA-seq marker genes. The “Pan Astrocyte” signature reflects top marker genes distinguishing all astrocytes from other non-neoplastic cell types. The “Astrocyte TE,” “Astrocyte NT,” and “Astrocyte R” signatures represent top marker genes specific to each astrocyte subcluster, identified using differential expression within the astrocyte lineage. Spatial module scores were computed on 16 μm Visium HD bins. Healthy brain tissue shows regionally distinct enrichment of TE and NT astrocyte signatures, whereas tumor sections display widespread expression of the reactive astrocyte signature, particularly at ventricular zones and tumor borders. **(D)** Dot plot showing expression of top marker genes for each astrocyte subcluster. Rows correspond to Astrocyte TE, Astrocyte NT, and Astrocyte R subclusters; columns show the top 10 marker genes for pan astrocytes and for each astrocyte subtype, identified by differential expression analysis. Dot color represents average normalized expression and dot size represents the percentage of cells expressing each gene within a given subcluster. **(E)** UMAP projection of all single cells, coloured by normalized Gfap expression (hexagon binning, median per bin). High Gfap expression is concentrated in the astrocyte manifold, with strongest signal in the reactive astrocyte subcluster. **(F and G)** Hallmark pathway activity analysis of astrocyte populations computed using AUCell. **(F)** Waterfall plot showing the top differentially enriched Hallmark gene sets distinguishing reactive astrocytes from TE/NT astrocytes, ranked by t-score. **(G)** Waterfall plot showing Hallmark gene set differences between reactive astrocytes from tumor tissue and those from healthy brain, similarly ranked by t-score. **(H)** Differential gene expression between reactive astrocytes from tumor tissue and healthy brain. The volcano plot highlights transcripts with significant log fold-change differences, with genes among the top upregulated genes annotated for reference. **(I)** Module scores for the reactive-astrocyte gene signature (top 10 subtype-specific markers) calculated across astrocytes from healthy, primary tumor, and recurrent tumor samples. Boxplots display the distribution of signature scores in each condition, with statistical comparisons performed using the Kruskal–Wallis test and pairwise Wilcoxon tests.

The third subcluster represented the dominant astrocyte population in tumor tissues. These cells expressed, in addition to the core astrocyte genes mentioned above, hallmark genes of the reactive astrocyte phenotype, including structural components (*Gfap*, *Vim*, *Cd44*, *Sulf2*)^23,25,26^ and neuroinflammatory or stress-induced transcripts (*C4b*, *Cebpd*, *S100a11*, *Ifitm3*, *S1pr3*)^27,28^ (Figure 3D–E). Spatial mapping of these genes showed enrichment throughout the tumor core but most prominently along perivascular regions, fiber tracts, ventricular zones, and tumor borders, consistent with glial-scar-like reactivity^29,30^ (Figure 3C). Expression of these genes was also detectable in healthy brain within periventricular and vascular niches, likely corresponding to subventricular-zone (SVZ) astrocytes (*Fabp7*, *Nes*, *S100a6*)^31–34^, which share portions of the reactive transcriptional program (Figure 3C and Supplementary Figure 3A).

Gene set enrichment analyses supported the reactive phenotype of these cells relative to TE/NT astrocytes, showing significant upregulation of pathways involved in apoptosis, IL-6 and complement signaling, coagulation, epithelial–mesenchymal transition (EMT), and TGF-β signaling (Figure 3G). Likewise, direct comparison of reactive astrocytes from tumor tissue versus their counterparts in healthy brain revealed strong enrichment of EMT, TNF-alpha signaling, hypoxia, p53, apoptosis, complement, and inflammatory response pathways (Figure 3H–I). Among the most induced genes were *Osmr*, *Igfbp2*, *Cebpd* and *S1pr3*, all associated with inflammatory activation, and *Ncan, Sdc3*, C*cn1 and Hbegf* linked to extracellular matrix remodeling and glial scar formation. A quantitative increase in the expression of reactivity-associated genes was evident when stratifying the reactive astrocyte cluster by sample type, with a marked elevation in tumor tissue and a further increase in recurrent post-irradiated samples (Figure 3I). Together, these findings demonstrate that tumor-associated astrocytes undergo a profound transcriptional reprogramming toward a highly reactive, cytokine- and ECM-driven state, likely contributing to tumor progression and recurrence, with this phenotype being most pronounced in post-radiotherapy recurrent tumors.

### Neoplastic cell heterogeneity in murine glioblastoma

The neoplastic cells were divided into ten transcriptionally distinct clusters within the original SNN graph-based clustering (Figure 2A and 4A). All ten neoplastic clusters exhibited high expression of the RCAS transgenes together with broad pan-neoplastic gene programs (Figure 4B–C). Three clusters (*Neopl-CC-I*, *-II*, and *-III*) expressed large sets of cell cycle–associated genes, including *Mcm3*, *Hist1h1b*, *Cenpa*, and *Cdc20* (Figure 4B), and were highly correlated with one another (Figure 4D). These clusters showed strong enrichment of G1/S and G2/M phase signatures derived from human GBM single-cell data^35^, indicating that they represent actively cycling GBM cell populations (Figure 4E). The proportion of cycling cells increased in recurrent tumor samples (Figure 4F), which was further confirmed by immunostaining for proliferation markers (Figure 4G).

**Figure 4.**
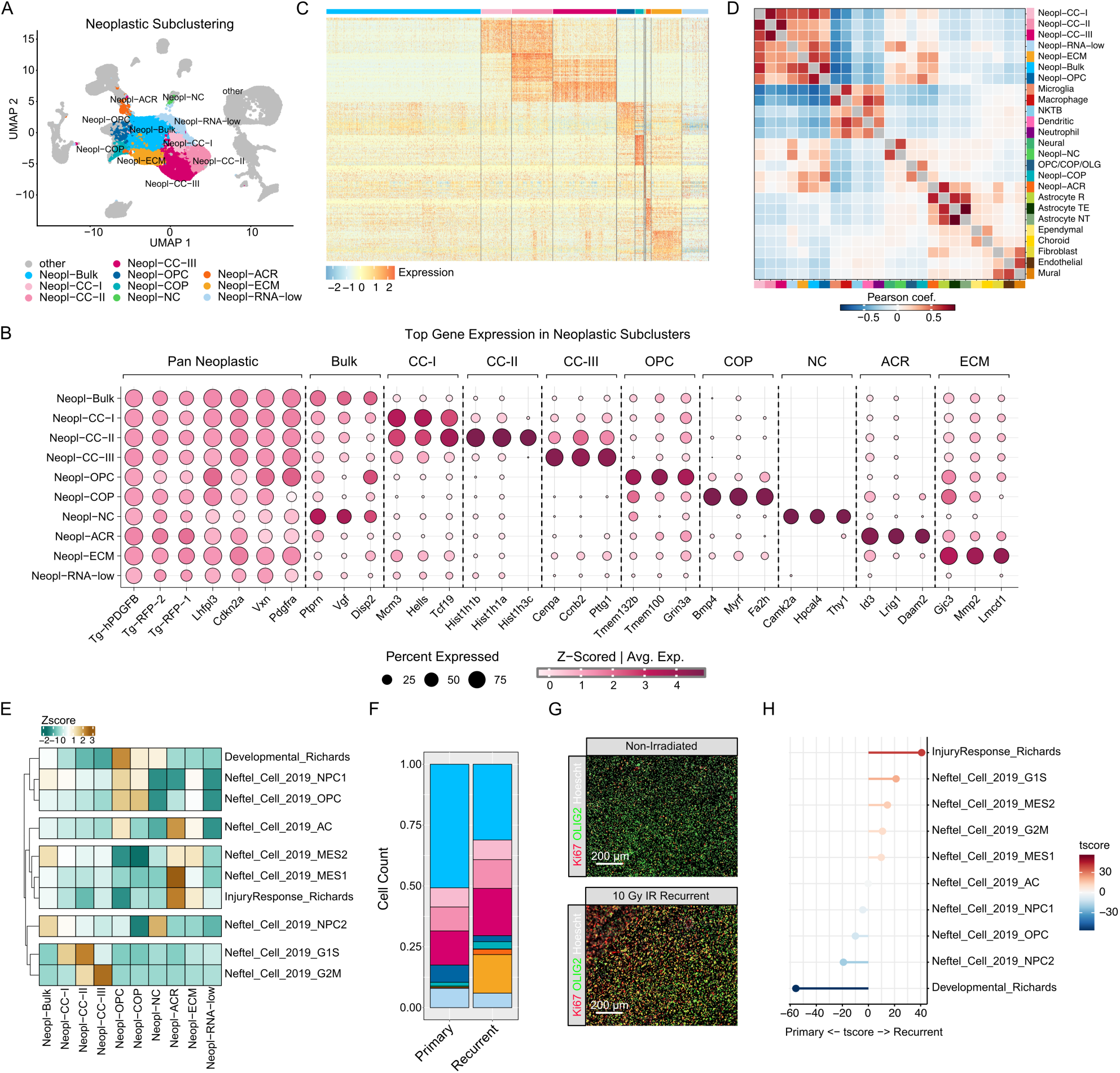
Neoplastic cell heterogeneity and lineage programs in PDGFB-driven murine GBM. **(A)** UMAP projection showing the ten neoplastic subclusters identified within the global SNN graph (same embedding as Figure 2A). Clusters correspond to distinct tumor cell states, including cycling populations (Neopl-CC-I, -CC-II, -CC-III), OPC-like and COP-like states, neural-like (Neopl-NC), reactive astrocyte–like (Neopl-ACR), ECM-associated (Neopl-ECM), a bulk population (Neopl-Bulk), and a low-RNA–content cluster (Neopl-RNA-low). Dot plot showing the top three marker genes defining each neoplastic subcluster. Dot size represents the percentage of cells expressing the gene, and color intensity indicates average scaled expression. **(C)** Heatmap displaying scaled expression levels of the top 50 marker genes for each neoplastic subcluster. Columns represent individual cells ordered by subcluster identity, and rows represent cluster-enriched genes. **(D)** Pearson correlation matrix comparing average gene-expression profiles of all neoplastic subclusters with annotated non-neoplastic cell types. Cycling clusters show strong mutual similarity, while non-cycling neoplastic clusters dider in their resemblance to OPC, COP, neuronal, or reactive astrocyte populations. **(E)** Heatmap of AUCell gene-set activity scores (z-scored) for Neftel and Richards GBM programs across neoplastic clusters. Rows show individual human GBM signatures and columns the ten neoplastic clusters; values are z-scores of AUCell enrichment statistics summarised per cluster and grouped into five signature modules. **(F)** Representative immunofluorescence images of primary (non-irradiated) and recurrent (10 Gy) tumors stained for Ki67 (proliferation), Olig2 (oligodendroglial/neoplastic marker), and Hoechst (nuclei), illustrating increased proliferative activity in recurrent tumors. Scale bar, 200 μm. **(G)** Relative abundance of neoplastic clusters in primary versus recurrent tumors. Stacked bar plots show the proportional contribution of each neoplastic state within each condition, highlighting decreased Neopl-Bulk/Neopl-OPC/Neopl-NC and increased Neopl-ACR, Neopl-ECM, and cycling (Neopl-CC) populations in recurrent tumors. **(H)** Waterfall plot of human GBM signatures gene-set activity in neoplastic cells comparing recurrent to primary tumors. Injury-response, mesenchymal, and cell-cycle programs are preferentially enriched in recurrent tumors, whereas NPC/developmental signatures are relatively higher in primary tumors.

Among the remaining neoplastic clusters, we observed broad variation in lineage-associated transcriptional programs corresponding to glial and neural cell states (Figure 4B and 4C). Correlation analyses with non-neoplastic reference populations revealed signatures resembling OPC, committed oligodendrocyte precursor (COP), neural, and reactive astrocyte–like profiles, illustrating that this murine model recapitulates the lineage plasticity characteristic of human GBM (Figure 4D). The largest cluster, designated Neopl-Bulk, showed the closest transcriptional similarity to the Neopl-OPC cluster and scored highest for the NPC2-like program from Neftel et al. (Figure 4E), consistent with a proneural, OPC/NPC progenitor-like phenotype.

Importantly, we observed that the proportions of Neopl-Bulk, Neopl-OPC, and Neopl-NC clusters decreased in recurrent tumors (Figure 4F and Supplementary Figure 4A). In contrast, the Neopl-ACR and Neopl-ECM clusters were almost exclusively present in recurrent samples. These clusters displayed strong associations with the AC/MES1/MES2 and Injury Response signatures described by Neftel *et al*^35^. and Richards *et al*.^36^, respectively. This pattern mirrors the cellular heterogeneity frequently observed in human GBMs, where tumor progression is accompanied by a shift from a proneural-like to a mesenchymal-like state^5,37–39^. Pathway enrichment analyses further substantiated this, revealing a significant upregulation of injury-response, mesenchymal, and cell-cycle programs in recurrent tumors, accompanied by reduced NPC2 and developmental/GSC program activity. (Figure 4H).

### Radiotherapy enriches glioma cells with a reactive astrocyte–like state

Analyses of both neoplastic and non-neoplastic compartments indicated that the tumorigenic process promotes a reactive astrocyte phenotype (Figure 3), and that this transcriptional program is not confined to astrocytes *per se* but can also be adopted by neoplastic cells, here termed neoplastic ACR cells. These neoplastic ACR cells displayed strong transcriptional similarity to non-neoplastic reactive astrocytes, as reflected by correlation analyses (Figure 4D) and their proximity in UMAP space (Figure 5A).

**Figure 5.**
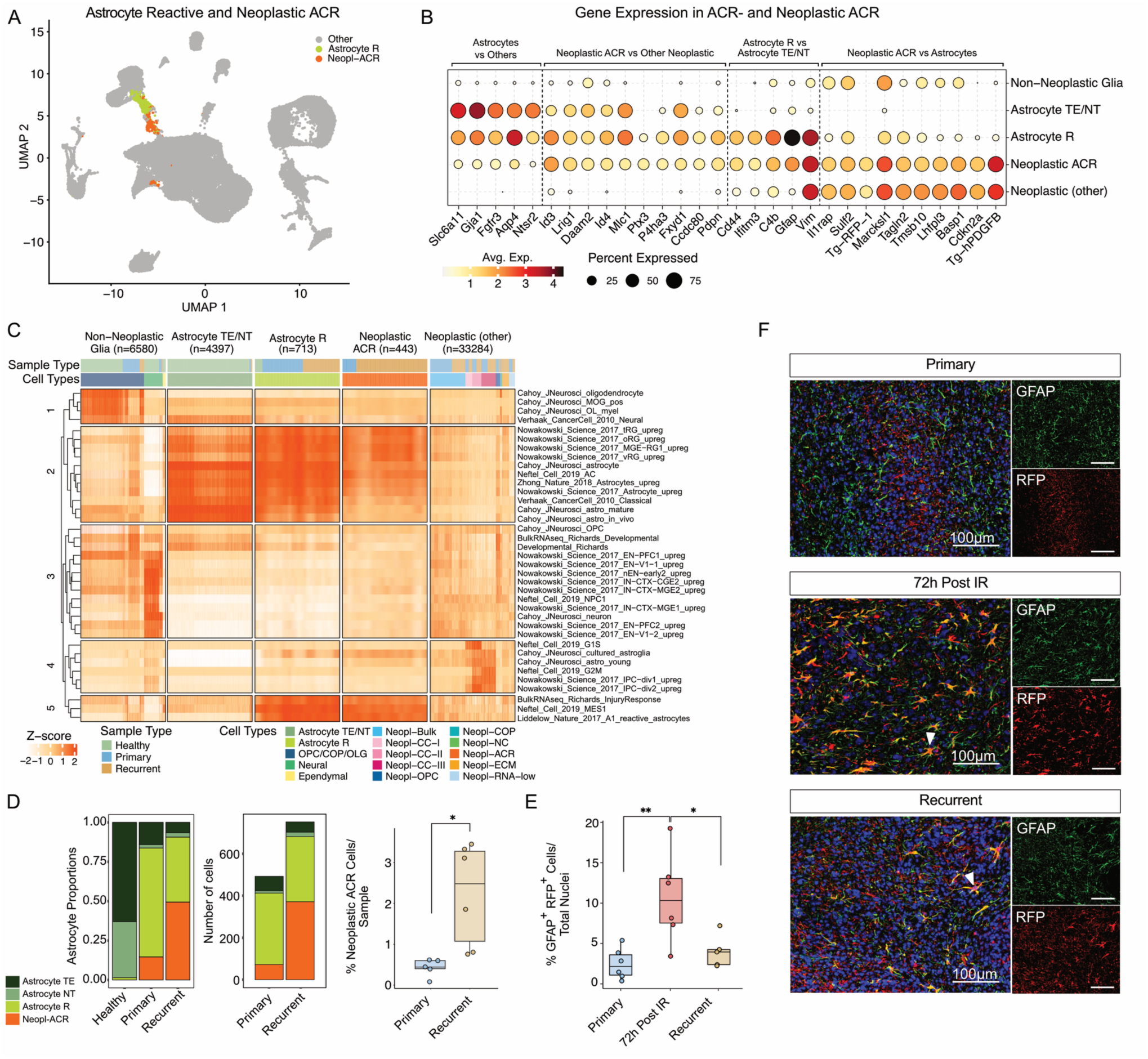
Radiotherapy enriches glioma cells with a reactive astrocyte–like phenotype. **(A**) UMAP projection of all 64,804 single cells highlighting non-neoplastic reactive astrocytes (Astrocyte R, green) and neoplastic ACR cells (orange); all other cells are shown in grey. **(B)** Dot plot summarizing expression of genes associated with astrocyte identity, reactive astrocyte activation, and neoplastic programs across five groups: non-neoplastic glia (other), Astrocyte TE/NT, Astrocyte R, Neoplastic ACR, and other neoplastic cells. Genes were selected from differential expression contrasts between astrocytes and other non-neoplastic cells, Astrocyte R versus TE/NT, Neopl-ACR versus other neoplastic cells, and Neopl-ACR versus astrocytes. Dot colour encodes average normalized expression and dot size the percentage of cells expressing each gene within a group, illustrating that Neopl-ACR cells co-express reactive astrocyte markers together with neoplastic signatures. **(C)** Heatmap showing AUCell-based pathway activity scores for curated neural and GBM gene signatures (Richards *et al*. 2021). Columns represent individual cells ordered within five major annotation groups: Non-Neoplastic Glia, Astrocyte TE/NT, Astrocyte R, Neoplastic ACR, and Neoplastic (other). Rows correspond to the top enriched signatures selected within each group (ranked by p value). Values represent row-wise z-scores. Reactive astrocyte, injury-response, and mesenchymal signatures are enriched in both Astrocyte R and Neoplastic ACR groups, whereas developmental and lineage-specific neuronal and OPC signatures dominate other compartments. **(D)** Distribution of astrocyte and neoplastic ACR populations across conditions. Left: relative proportions of Astrocyte TE, Astrocyte NT, Astrocyte R, and Neoplastic ACR cells across healthy, primary, and recurrent tumors. Middle: absolute counts of astrocyte and neoplastic ACR cells per tumor type. Right: percentage of Neoplastic ACR cells per sample, showing a significant increase in recurrent tumors. **(E)** Quantification of GFAP and RFP co-positive cells in primary tumors, tumors collected 72 hours after irradiation, and recurrent tumors. Boxplots show the percentage of GFAP+ RFP+ cells over total nuclei per sample. Statistical comparisons were performed using the Kruskal–Wallis test with pairwise Wilcoxon tests. **(F)** Immunofluorescence staining for GFAP (green), RFP (red), and DAPI (blue) in primary, early post-irradiated (72 h), and recurrent tumors. RFP identifies tumor-derived cells, and arrows indicate GFAP+ RFP+ double-positive tumor cells. Acute induction of GFAP in tumor cells is evident 72 h after irradiation and persists in recurrent lesions (One-way ANOVA, Tukey’s multiple comparison test). Scale bars: 100 μm.

When comparing gene expression of neoplastic ACR cells with non-neoplastic astrocyte subclusters and other neoplastic populations, we observed co-expression of canonical neoplastic genes together with a broad panel of reactive astrocyte markers, including *Gfap*, *Vim*, *Cd44*, and *C4b*, as well as extracellular-matrix and secreted factors implicated in glial scar formation such as *Ptx3*, *Fxyd1*, and *Pdpn*^27,28^ (Figure 5B).

Pathway analysis using a broad range of curated brain and GBM cell-type signatures revealed that neoplastic ACR cells exhibited high enrichment of reactive astrocyte, “injury-response”^36^, and mesenchymal-like^35^ programs, alongside increased scores for astrocytic and astrocyte-progenitor signatures (Nowakowski *et al.*^40^ signatures for truncated radial glia (tRG), outer radial glia (oRG), ventricular radial glia (vRG)) (Figure 5C). These increases were accompanied by a relative reduction of mature astrocyte pathways, consistent with a shift toward a reactive or progenitor-like state.

At the cellular level, reactive astrocytes were present in both primary and recurrent tumors, whereas neoplastic ACR cells were markedly enriched in established recurrent lesions (Figure 5D). To determine whether this phenotype emerges acutely following irradiation, we analyzed tumors 72 h post-radiotherapy using RFP and GFAP co-staining to identify tumor-derived astrocyte-like cells. We detected a significant increase in RFP⁺/GFAP⁺ tumor at 72 h post-radiotherapy, as compared to untreated primary tumors (Figure 5E–F). Together, these findings demonstrate that radiotherapy rapidly induces or selects for tumor cells with a reactive astrocyte-like phenotype, which persists in recurrent disease.

### Astrocyte-tumor cell communication during tumor progression

To investigate potential cell–cell communication within the glioblastoma microenvironment, we applied CellChat^41,42^ to infer ligand receptor interactions across all annotated cell types. Reactive astrocytes exhibited the strongest predicted outgoing interactions toward other astrocytes and neoplastic cells and were predicted to receive the majority of incoming interactions from fibroblasts and neoplastic cells (Figure 6A). Similarly, neoplastic cells displayed the highest number of predicted outgoing and incoming interactions with astrocytes and other neoplastic cells (Figure 6A).

**Figure 6.**
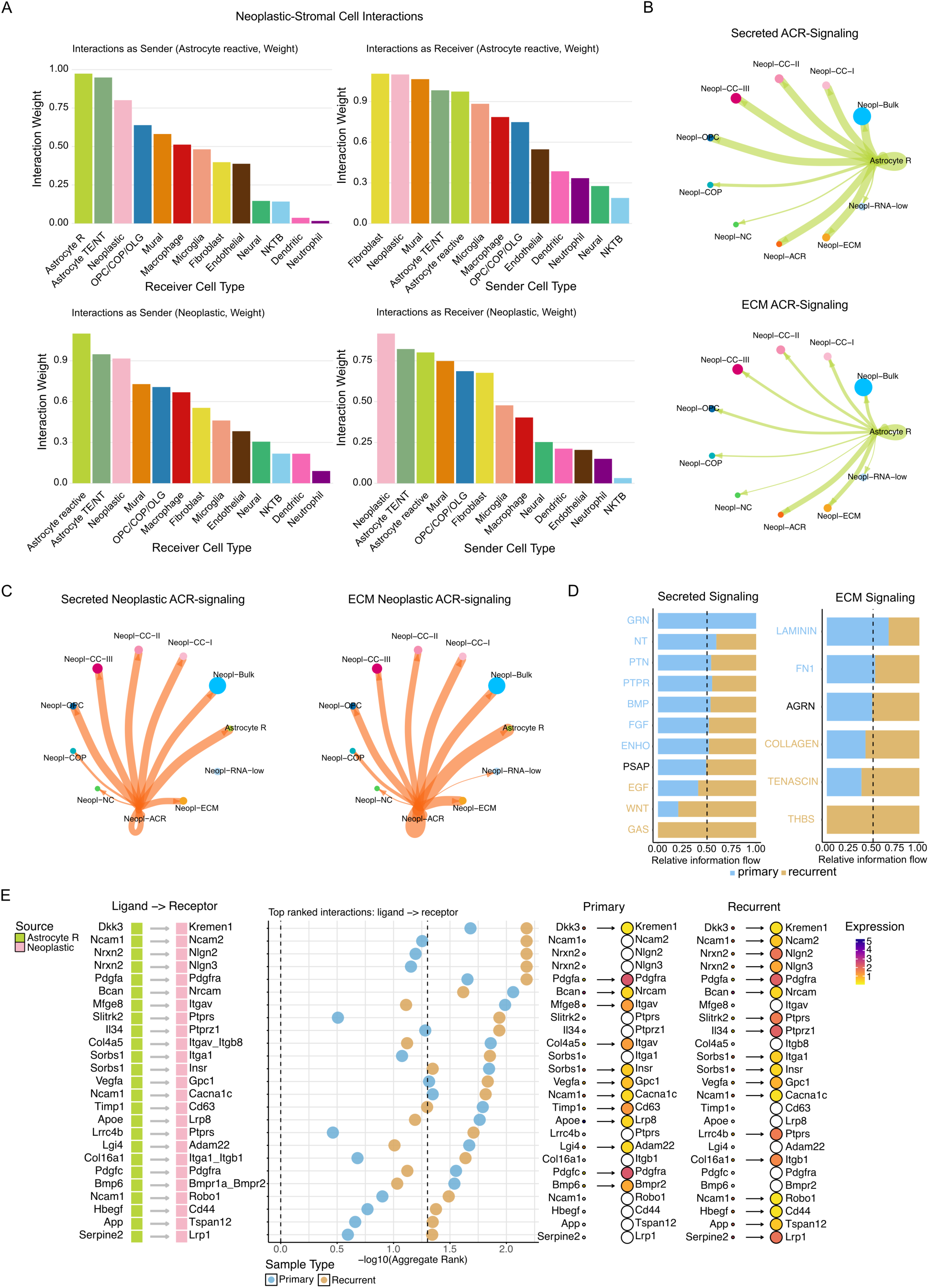
Astrocyte–tumor cell communication in primary and recurrent PDGFB-driven GBM. **(A)** CellChat analysis of secreted signaling between major cell populations. Bar plots show summed interaction weights for reactive astrocytes (Astrocyte R, top row) and neoplastic cells (bottom row) acting as sender (right) or receiver (left). Bars are ordered by interaction weight, and colors indicate the interacting cell types. **(B)** Circle network plots of CellChat-inferred signaling from reactive astrocytes to the ten different neoplastic subclusters. Top: secreted signaling. Bottom: ECM–receptor signalling. Edge thickness reflects interaction weight and node size reflects the number of cells per subcluster. **(C)** Comparison of pathway-level predicted signaling from reactive astrocytes to neoplastic cells between primary and recurrent tumors. Bar plots show relative information flow for selected secreted (left) and ECM-related (right) signaling pathways, with blue and brown bars indicating primary and recurrent tumors, respectively; the dashed line marks equal contribution of both conditions. **(D)** Circle network plots of Neopl-ACR-derived signaling to neoplastic subclusters and reactive astrocytes. Left: secreted signaling. Right: ECM–receptor signaling. Edge thickness and node size are proportional to interaction weight and cell number, respectively, highlighting preferential ECM-mediated communication between Neopl-ACR, Neopl-ECM, and Astrocyte R populations. **(E)** LIANA-based integrative ligand–receptor analysis comparing Astrocyte R → Neoplastic signaling between primary and recurrent tumors. The analysis identifies ligand–receptor pairs that are both statistically significant (aggregate rank < 0.05) and exhibit the largest differences in aggregate rank between conditions, thereby highlighting interactions with differential signaling patterns. Left: ligand (green) and receptor (pink) complexes contributing to the top differentially ranked interactions. Middle: dot plot showing the top-ranked ligand–receptor pairs, where the x-axis indicates –log10(aggregate rank), dot color indicates sample type (primary vs recurrent), and dot size reflects the number of target neoplastic cells. Right: per-condition summaries showing target-group size and the average expression levels of ligand and receptor genes in primary and recurrent tumors; arrows indicate whether an interaction passes the significance threshold (–log10(0.05)), corresponding to the dashed line shown in the middle panel.

We next examined predicted interactions from reactive astrocytes to the different subpopulations of neoplastic cells. Secreted signaling from reactive astrocytes was broadly distributed across neoplastic clusters, whereas extracellular matrix (ECM) mediated interactions were concentrated between reactive astrocytes and the neoplastic ACR and ECM subclusters (Figure 6B). Similarly, when analyzing predicted outbound interactions for the Neopl-ACR tumor subtype, a strong bias was observed for ECM associated signaling toward reactive astrocytes and the Neopl-ECM subtype (Figure 6C). These results indicate predicted preferential and reciprocal communication between reactive astrocytes and neoplastic ACR cells, with ECM remodeling as a dominant mode of interaction.

We next assessed how predicted signaling from reactive astrocytes to neoplastic cells differed between primary and recurrent tumors. Several pathways were selectively enriched in recurrent tumors, including GAS, WNT, EGF, THBS, Tenascin, and collagen signaling (Figure 6D), implicating both extracellular and growth factor signaling in the recurrent tumor microenvironment^43–47^. Blocking the Gas6/Axl-axis in PDGFB-induced glioma primary cultures (PIGPC) resulted in reduced proliferation and colony formation capability (Supplementary Figure 5A, 5D), whereas culture with HB-EGF and Tenascin C significantly increased PIGPC colony formation with or without 4 Gy radiotherapy without significantly affecting tumor cell proliferation (Supplementary Figure 5B-C, 5E-F). To confirm co-localization of reactive astrocytes and Tenascin C in murine GBM, we performed immunofluorescent staining of Tenascin C with GFAP and Olig2, showing Tenascin C expression in tumor areas with high GFAP expression (Supplementary Figure 5G). These findings suggest that reactive astrocyte-derived *Gas6*, *Hbegf* and *Tnc* may support tumor progression. To further identify ligand-receptor interactions that differ between primary and recurrent tumors, we applied the LIANA framework, which integrates and harmonizes results from multiple published inference methods and ligand receptor databases (Figure 6E). This analysis revealed recurrence-specific enrichment of interactions involved in cell adhesion and synaptic regulation (*Ncam1, Nrxn2*, *Slitrk2*, *Lrrc4b*, *App*), extracellular matrix remodeling and tissue repair (*Col16a1*, *Vegfa*, *Serpine2*, *Hbegf*), and immune microenvironment modulation (*Il34*).

Together, these findings suggest that the reactive astrocyte phenotype is an important contributor to the cellular communication network of the GBM microenvironment and that astrocyte–tumor cell crosstalk is particularly enhanced in the post-radiotherapy recurrent setting through extracellular matrix and growth factor mediated pathways.

### Reactive astrocyte–like tumor states emerge and expand during GBM progression and recurrence

We next applied gene signatures derived from the scRNA-seq dataset to predict the spatial distribution of neoplastic and non-neoplastic populations in the Visium HD sections. Spots located within the tumor core were predominantly assigned to OPC-like and bulk-like neoplastic states, with these regions bordered by bands enriched in cycling tumor cells. Notably, Neopl-ACR predictions localized most frequently to regions immediately adjacent to areas mapped as reactive astrocytes, including ventricular zones and peripheral regions at the tumor margin (Figure 7A; see also Figure 3B). Co-localization of tumor-derived and non-neoplastic GFAP-positive cells was further supported by multiplex immunofluorescence, which revealed both RFP⁺/GFAP⁺ and RFP⁻/GFAP⁺ populations within peritumoral niches (Figure 7B).

**Figure 7.**
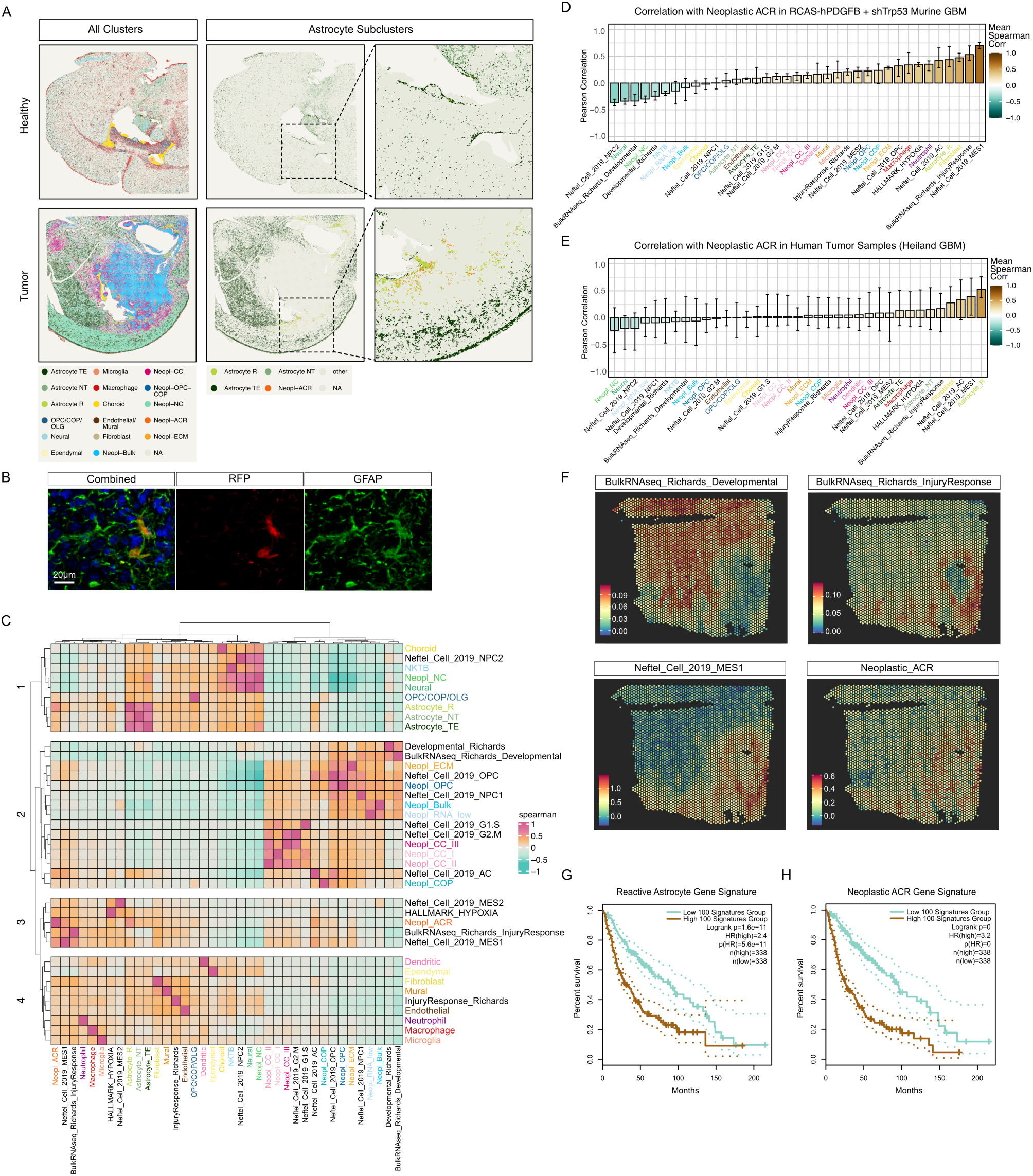
Reactive astrocyte–like tumor states emerge and expand during GBM progression and recurrence. **(A)** Spatial mapping of scRNA-seq–derived cell states onto Visium HD sections using Seurat transfer anchors. Left: predicted major cell populations for a healthy brain section (top) and a tumor-bearing section (bottom). Middle and right: corresponding maps showing only astrocyte and Neopl-ACR subclusters, with zoom-in views highlighting regions around the lateral ventricle and tumor margin. Note the close apposition of Neopl-ACR spots (orange) to regions enriched for reactive astrocytes (yellow). **(B)** Representative immunofluorescence image of a tumor section showing RFP-labelled tumor cells (red) and GFAP-positive astrocytes (green) with nuclear counterstain (blue). Both RFP-positive and RFP-negative GFAP^+^ cells are observed in peritumoral regions. Scale bar, 20 μm. **(C)** Spearman correlation heatmap of module scores for scRNA-seq–derived mouse programs and human GBM gene signatures calculated from spatial RNA-seq Visium HD data from a tumor sample. Modules group into four major clusters (1– 4), with strong positive correlations between Neopl-ACR, reactive astrocyte, Neftel MES1, hypoxia (HALLMARK_HYPOXIA), and the Richards injury-response signatures. **(D)** Ranked mean Spearman correlations between the Neopl-ACR module score and all other signatures across four in-house Visium v1 samples. Bars show the mean correlation across the four samples; error bars indicate the minimum and maximum correlation observed. **(E)** As in (D), but for human GBM Visium v1 data from Heiland *et al*., restricted to tumor samples. Neopl-ACR correlations are consistently highest with MES1-like, hypoxia-associated, and injury-response programs. **(F)** Representative GBM Visium v1 sections from Heiland (2019) study showing spatial module scores for the Richards developmental and injury-response astrocyte signatures, Neftel MES1, and Neopl-ACR. Neopl-ACR and MES1 scores are concentrated in hypoxic or stressed tumor regions that overlap with high injury-response astrocyte scores. **(G,H)** Kaplan–Meier overall-survival analyses in the TCGA cohort using the reactive astrocyte (G) and Neopl-ACR (H) signatures. Patients were stratified into high- and low-score groups (top and bottom 100 cases, respectively). Both signatures are significantly associated with reduced overall survival (log-rank test p values and hazard ratios indicated in the panels).

We then quantified module scores for a combined set of scRNA-seq derived programs and curated human GBM signatures (Figure 7C). Spatial variation in these signatures recapitulated the transcriptional relationships observed in the single-cell data (Figures 4E and 5C). Neopl-ACR module scores correlated strongly with reactive astrocytes, and showed an even stronger correspondence with the Richards injury-response program^36^ and the MES1-like state described by Neftel *et al*.^35^. Additional associations were observed with hypoxia-related genes and the MES2 program. Module scores for these signatures displayed overlapping spatial distributions, with Neopl-ACR, MES1, and injury-response scores co-enriched in hypoxic or otherwise stressed regions of the tumor microenvironment (Figure 7C).

These associations were also evident in Visium v1 data from two primary and two post-radiotherapy recurrent tumors (Figure 7D and Supplementary Figure 6A), and were further replicated in an external human spatial dataset comprising twenty GBM samples^14^ (Figure 7E-F and Supplementary Figure 6B). To assess clinical relevance, we applied the reactive astrocyte and Neopl-ACR signatures to the TCGA cohort of low- and high-grade gliomas. Both signatures were significantly associated with decreased overall survival (Figure 7G-H). Taken together, the spatial and transcriptional patterns show that reactive astrocyte and Neopl-ACR states correlate with adverse tumor characteristics in both the murine model and human GBM samples.

## Discussion

Intratumoral heterogeneity and phenotypic plasticity are increasingly recognized as key drivers of aggressive growth and therapeutic resistance in GBM^48,49^. Such diversity extends beyond the neoplastic compartment to include multiple cell types within the tumor microenvironment. While genetic cues can influence which cell states predominate, accumulating evidence suggests that local environmental conditions within distinct tumor niches also promote state transitions among neoplastic cells. Here, we sought to determine how stress associated with tumor initiation, progression, and particularly radiotherapy, shapes phenotypic changes in both neoplastic and non-neoplastic cells in GBM. We observed marked alterations in cellular composition in post-radiotherapy recurrent tumors, including a previously described shift from microglia-dominant primary tumors to macrophage-dominant recurrent tumors^50^. Concurrently, neoplastic cells in recurrent tumors exhibited transcriptional and phenotypic features characteristic of reactive astrocytes, including activation of ECM remodeling programs. These reactive-like tumor states were closely aligned with those observed in non-neoplastic tumor-associated astrocytes during tumor formation. Together, these findings support a model of coevolution between tumor and stromal cells under stress and underscore the role of the microenvironment in shaping the recurrent tumor landscape.

Recent single-cell and spatial transcriptomic studies have positioned GBM cells along a continuum between developmental and injury-response states^36,51,52^. Consistent with this framework, we observed a progressive shift of neoplastic cells toward injury-response programs as tumors evolved from primary to post-radiotherapy recurrence in our murine model. Our data support a model in which injury-response signals from both tumor and stromal compartments cooperate to drive this phenotypic plasticity. In line with this, cell-cell communication inference analyses revealed that reactive astrocytes are predicted to engage in close signaling interactions with neoplastic ACR-like cells specifically in recurrent tumors. The majority of these interactions appear to involve ECM remodeling, consistent with recent reports describing fibrosis-like scar formation in post-radiotherapy GBM^53,54^. These observations further implicate reactive astrocytes as central architects of the recurrent GBM microenvironment, promoting tumor persistence and regrowth^55^.

Previous studies have linked GBM progression to a phenotypic shift toward mesenchymal (MES)-like states^56–60^, which are frequently associated with more aggressive and therapy-resistant behavior^61,62^. For instance, resistance in primary human GBM cultures has been correlated with MES-like signatures that share molecular similarities with reactive astrocyte programs^36,63,64^. In agreement with this, we found that the neoplastic ACR state emerging in recurrent murine GBM closely resembles both the MES1 state defined by Neftel *et al*. and the injury-response state described by Richards *et al*. Although we did not directly assess the relationship between this cell state and therapeutic resistance, we observed a dramatic increase in Gfap⁺RFP⁺ neoplastic ACR cells shortly after radiotherapy, that partially persisted at the time of recurrence. This may reflect either selection for inherently resistant cells or rapid induction of the ACR-like state in response to radiotherapy-induced stress. Importantly, both reactive astrocyte and neoplastic ACR transcriptional signatures were associated with aggressive disease and poor outcome in human GBM, further supporting their relevance to recurrence.

Notably, not all longitudinal human GBM studies have identified consistent MES shifts between primary and recurrent tumors^65^. This discrepancy likely reflects sampling biases inherent to clinical biopsies, often limited to selected regions, compared to the more comprehensive whole-tumor analyses possible in murine models. Additionally, the extensive genetic heterogeneity of patient tumors may obscure subtle or variable shifts that are more readily detectable in genetically uniform experimental models. Nonetheless, independent systems, including human tissue–based models, converge on the conclusion that tumor cells tend to adopt MES-like or injury-response phenotypes upon recurrence.

In conclusion, our findings support a model in which reciprocal interactions between reactive glia and tumor cells drive phenotypic plasticity and therapeutic adaptation, ultimately shaping the trajectory of GBM recurrence. Understanding how reactive glial signaling fuels tumor cell plasticity may reveal new therapeutic opportunities to disrupt the adaptive programs that underlie GBM recurrence.

## Acknowledgements

Funded by the European Union (ERC, RESISTANCEPROGRAMS, 101043587). Views and opinions expressed are however those of the author(s) only and do not necessarily reflect those of the European Union or the European Research Council Executive Agency. Neither the European Union nor the granting authority can be held responsible for them. This study was further supported by the Swedish Cancer Society, the Swedish Research Council, the Swedish Childhood Cancer Fund, Ollie & Elof Ericssons foundation, and the Crafoord foundation. We gratefully acknowledge the support by Mrs. Berta Kamprad’s Cancer Foundation to the L2CancerBridge program at CREATE Health Cancer Center.

## Material and Methods

### Generation of Murine Gliomas

Murine gliomas were generated using Nestin/tv-a (Ntv-a) mice, by injecting RCAS-shTrp53 and RCAS-hPDGFB-transfected DF-1 cells (ATCC ® CRL-12203™, ATCC, Manassas, VA, USA) intracranially into the neonatal mouse brain, as previously described^18,19^. Mice were observed daily and euthanized upon glioma symptoms. For the transcriptomics analyses, radiotherapy was performed around the time of glioma development, on days 31-35. Cranial radiotherapy was administered through a single fraction of 10 Gy using a preclinical research platform (XenX, XStrahl Inc, Suwanee, GA, USA). For short-term irradiation experiments, radiotherapy was administered at day 42 post-injection, and mice were harvested 72 hours later.

Mice were sacrificed upon exhibiting glioma symptoms. Tumors or corresponding brain regions from healthy tissue were dissected. All tissue processing was performed on ice. Tumor samples intended for 10x Spatial Transcriptomics were immediately snap-frozen in isopentane on dry ice, while the remaining tissues were placed in pre-chilled PBS on ice.

### Multiplex Immunofluorescence

Whole brain from euthanized animals embedded in Optimal Cutting Temperature Compound (OCT) (Thermo Fisher Scientific) were snap-frozen in ice-cold isopentane. Sections were dried at room temperature for 30 minutes, followed by fixation in ice-cold acetone. Sections were permeabilized with 0.3% Triton X-100 in PBS followed by blocking in 3% BSA. Primary antibody incubation was performed in 3% BSA for 1 hour at room temperature, or overnight at 4°C.

To evaluate tumor cell presence, we performed multiplex immunofluorescence with an antibody against RFP, which is expressed from the shTrp53-encoding RCAS vector. Primary antibody staining was used targeting RFP (Abcam, ab185921, 1:50 dilution), Olig2 (R&D systems, AF2418, 1:200 dilution), and GFAP (Abcam, ab4674, 1:400 dilution) or for the extracellular matrix staining with Tenascin C (Abcam, ab108930). After PBS wash, the sections were incubated with appropriate secondary AlexaFluor-conjugated antibodies for 1 hour in room temperature. Cell nuclei were marked with 4′,6-diamidino-2-phenylindole (DAPI) (Sigma-Aldrich). To control for nonspecific signals, tissue sections were incubated with secondary antibodies alone. Scanning of slides was performed on PhenoImager (Akoya Biosciences). A spectral fluorophore for each primary antibody was used to create a library and perform spectral unmixing of images using the InForm software (Akoya Biosciences). Image analysis was performed in QuPath. Cell detection was based on DAPI staining, and multiplexed analysis was performed using Machine learning on training images according to QuPath tutorials (https://qupath.readthedocs.io)^66^.

### Isolation of tumor cells from mouse

Tumor cells were collected from a sacrificed mouse with glioma symptoms by dissecting out tumor tissue. Single cell suspension was performed using the Adult Brain Dissociation Kit for mouse and rat tissue (Merck #130-107-677) and were resuspended in FACS buffer. Cells were stained with BD Via-Probe™ Red Nucleic Acid Stain (5 nM), and analyzed by flow cytometry. Live cell sorting was performed based on live cells with clear RFP positivity on BD FACSMelody™ Cell Sorter. Cells were cultured in serum-free HGC medium on poly-ornithine and laminin-coated plates and passaged using Accutase (Thermo Fisher Scientific). Cells were incubated in a 37°C humidified incubator with 21% O_2_ and 5% CO_2_.

### Proliferation assay

Cells were seeded at a density of 5000 cells/well in a 96-well plate. Day 2, cells were treated with one of the following recombinant proteins; 100ng/ml Gas6 (R&D Systems, 986-GS-025/CF), 5μg/ml Tenascin C (R&D Systems, 3358-TC-050) or 25ng/mL HB-EGF (R&D Systems, 259-HE-050/CF) or 100nM, 500nM or 1μM Bemcentinib (R428), a selective Axl-inhibitor. For investigation with Gas6 and R428, 96-well plates were coated with poly-orthinine and laminin before cell seeding and cells were additionally treated with or without 4 Gy radiotherapy using a Cix1 x-ray irradiator (Xstrahl) 1 hours after addition of recombinant protein and Axl-inhibitor. Cell viability was measured at 24, 72 and 120 hours with addition of WST-1 (Abcam, ab155902) to the medium. Absorbance was measured at 450 nm in Synergy 2 Plate reader (BioTek).

### Colony formation assay

PIGPC cells were plated at 250 cells/well in 6-well plate. On day 2, cells were treated with one of the following recombinant proteins; 100ng/ml Gas6 (R&D Systems, 986-GS-025/CF), 1μg/ml Tenascin C (R&D Systems, 3358-TC-050) or 25ng/mL HB-EGF (R&D Systems, 259-HE-050/CF) or 100nM Bemcentinib (R428) for blocking Gas6-Axl-signaling. For investigation with Gas6 and R428, 96-well plates were coated with poly-ornithine and laminin prior to cell seeding. All cells were then treated with or without radiotherapy at 4 Gy using a Cix1 x-ray irradiator (Xstrahl). PIGPC cells were then incubated until visible colony formation for up to two weeks. Following fixation with 4% PFA, colonies were stained with 0.01% Crystal Violet for 1 hour, then imaged on the LAS-3000 system and quantified using ImageJ/Fiji (version 2.1.0) or counted by hand.

### Single-cell RNA sequencing (10x Genomics Flex)

Single-cell suspensions were generated from freshly dissected tumors and healthy brain tissue using the Adult Brain Dissociation Kit (Miltenyi Biotec 130-107-677). Suspensions with viability above 70% were processed using the Chromium Next GEM Single Cell Fixed RNA Sample Preparation Kit (10x Genomics 1000414) according to protocol CG000478. Fixation was performed by incubating cells in freshly prepared 4% formaldehyde in Conc. Fix & Perm Buffer (10x Genomics PN-2000517) for 24 hours at 4°C, followed by quenching and storage in 50% glycerol at –80°C until library preparation.

Fixation of Single-Cell Suspension was performed according to Protocol from Fixation of Cells & Nuclei from Chromium Fixed RNA Profiling (10x Genomics, CG000478) with Chromium Next GEM Single Cell Fixed RNA Sample Preparation Kit (16rxn) (10x Genomics, 1000414) by freshly preparing the fixation buffer containing 4% Formaldehyde diluted in Conc. Fix & Perm Buffer (10x Genomics, PN-2000517). Samples were fixed 24 hours at 4°C followed by Quenching and storing in 50% glycerol solution at -80°C until sequencing.

Whole-transcriptome libraries were generated using the Chromium Next GEM Single Cell Fixed RNA Flex kit (10x Genomics 1000568) at the Center for Translational Genomics, Lund University. Fixed cells were hybridized with the Chromium Mouse Transcriptome Probe Set v1.0.1 (10x Genomics). Cells were partitioned into Gel Bead–in–Emulsions (GEMs) on a Chromium Controller, and libraries were prepared according to the manufacturer’s instructions.

To detect cell specific RCAS-derived transgene expression, pilot experiments were performed with a set of candidate Flex probes targeting human PDGFB fused to HA (hPDGFB-HA), RFP, shTrp53, Tva, and GFP (negative control). Based on performance in these pilot runs, one hPDGFB probe and two RFP probes were selected for inclusion in the final panel; remaining probes targeted lowly expressed transcripts or showed poor performance and were not retained. Custom probe sequences are listed in Supplementary Table 1. In total, five Flex hybridization runs (including one technical replicate) yielded 23 libraries; two samples with near-zero UMI counts were excluded. Libraries were sequenced on an Illumina NovaSeq 6000 using an S4 Reagent Kit v1.5 (200 cycles).

### scRNA-seq preprocessing, quality control, and normalization

Raw FASTQ files were processed with Cell Ranger v7.1.0 (cellranger count) using a custom reference generated with cellranger mkref based on mm10 (refdata-gex-mm10-2020-A) with added transgene sequences. Initial quality-control metrics of 92,737 cells included UMI counts per cell (nCount), number of detected genes (nFeature), and the proportions of mitochondrial and haemoglobin reads. Cells with >10% mitochondrial or >5% haemoglobin reads were removed, and cells in the extreme top or bottom 5% of nCount and nFeature distributions were excluded. Putative doublets were identified and removed using scDblFinder^67^. To ensure high data integrity, four samples failing to meet stringent QC criteria were excluded. To ensure balanced representation of biological conditions, cells were subsequently downsampled to equalize group sizes across healthy brain, primary tumor, and recurrent tumor samples. This resulted in a final dataset comprising 64,804 cells: 22,655 cells from five primary tumors, 22,655 cells from six recurrent tumors, and 19,494 cells from five healthy samples.

To mitigate differences in sequencing depth across samples, UMI counts were downsampled to a maximum of 10,000 UMIs per cell using the Python package pynorm^68^. These UMI-normalized data were used for downstream dimensionality reduction, clustering, and visualization. All QC-filtered samples were merged into a single Seurat object.

### Dimensionality reduction, clustering, and cell type annotation

Downsampled UMI counts were normalized using Seurat’s NormalizeData function (log-normalization with a scale factor of 10,000). Highly variable genes (2,000–5,000) were identified using FindVariableFeatures, and data were scaled with ScaleData. Principal component analysis was performed with RunPCA using 30 principal components, and UMAP embeddings were computed from these components.

A shared nearest-neighbor (SNN) graph was constructed using FindNeighbors, and clustering was performed with FindClusters using the Louvain algorithm at a resolution of 0.5 unless otherwise indicated. All clustering was carried out on the global dataset; neoplastic cells were not re-clustered in isolation. Cell types were annotated using the Allen MapMyCells tool (RRID:SCR_024672) https://portal.brain-map.org/atlases-and-data/bkp/mapmycells in combination with curated marker gene sets.

To identify neoplastic cells, we constructed an RCAS-derived signature based on hPDGFB and RFP expression counts and used it to assign each cell as RCAS^high^, RCAS^low^, or RCAS^neg^ (Supplementary Figure 1). RCAS status was determined independently of clustering and subsequently mapped onto the SNN clusters. Mapping RCAS transgene expression onto the SNN clusters revealed three compartments: a large group containing only RCAS^neg^ cells, a mixed group containing both RCAS^neg^ and RCAS^low/high^ cells, and a neoplastic group composed almost exclusively of RCAS^low/high^ cells. We therefore classified RCAS^neg^ cells in the non-neoplastic and mixed groups as non-neoplastic, and all cells in the neoplastic group together with RCAS^low/high^ cells in the mixed group as neoplastic. Cells with discordant profiles were labelled ambiguous and excluded. This yielded 30,345 non-neoplastic cells, 33,727 neoplastic cells, and 732 ambiguous cells. Supplementary Figure 2 summarizes the cluster-level RCAS composition and sample-type distribution underlying this classification.

### Differential expression and module score computation

Differential gene expression between astrocyte subsets and between neoplastic clusters was performed using Seurat’s FindAllMarkers (Wilcoxon test, adjusted p-values). Cluster-specific marker sets were used to define astrocyte TE, NT, R signatures, neoplastic state signatures, and the RCAS module score. Gene signature scores were computed using Seurat AddModuleScore.

### Gene set enrichment and pathway scoring

Pathway enrichment was performed using AUCell via GeneSetAnalysis function in SeuratExtend package. AUCell scores were generated for curated sets of neural, glial, and glioma-associated signatures, including Hallmark Gene Ontology (MSigDB), Neftel *et al.*^35^ (AC, OPC, NPC1/2, MES1/2, G1/S, G2/M), Richards *et al*^36^. (injury-response and developmental programs), and additional astrocyte and lineage signatures from Cahoy *et al*.,^69^ Liddelow *et al*.,^70^ Nowakowski *et al.*,^40^ and Verhaak *et al*^71^. AUCell enrichment values were row-scaled and visualized with ComplexHeatmap.

### Cell–cell communication analysis

Cell–cell communication was inferred using CellChat^41,42^ applied to the QC-filtered, UMI-downsampled Seurat object (64,804 cells). Communication networks were computed using the CellChatDB.mouse database, run separately for Secreted Signalling and ECM–Receptor categories. Over-expressed genes and interactions were identified with default CellChat procedures, communication probabilities were computed using the triMean method with 1,000 bootstraps, and interactions supported by fewer than 25 cells per group were removed. Pathway communication probabilities were aggregated and used for downstream comparisons. Circle plots and summary matrices were generated from the CellChat interaction network to visualize outgoing and incoming signalling. To compare primary and recurrent tumors, CellChat models were run separately per condition, merged, and contrasted using rankNet, focusing on Astrocyte R-to-neoplastic signalling.

Complementary ligand–receptor inference was performed using LIANA on normalized RNA counts^72^. LIANA was run separately on primary, and recurrent tumor subsets, restricting analyses to Astrocyte R (sender) and Neoplastic cells (receiver). The multi-method mode was used, and consensus interaction scores were summarized as aggregate_rank, with lower rank indicating stronger support. For differential analyses, ligand–receptor pairs were harmonized across conditions and filtered for interactions with aggregate_rank < 0.05 in at least one condition. The top interactions showing the largest differences in –log10(aggregate_rank) between primary and recurrent tumors were selected for visualization. Mean expression of ligand and receptor genes in each sender/receiver group was computed, and results were displayed in combined dot-plot and summary panels indicating aggregate rank, target-group size, and per-condition expression, with significance referenced to the –log10(0.05) threshold.

### Visium HD Spatial Transcriptomics

One paraffin-embedded healthy mouse brain and one RCAS-derived tumor were processed using Visium HD Spatial Gene Expression chemistry (10x Genomics). Brains were post-fixed in 4% PFA for 24 hours, dehydrated, and embedded in paraffin. Sections (3 µm) were cut and mounted onto standard glass slides. For RNA quality control, ten 5 µm sections were processed with the RNeasy FFPE Kit (Qiagen 73504) and analyzed on a Bioanalyzer to ensure DV200 values above the recommended threshold.

Glass slides were H&E stained prior to probe hybridization. Hybridization was performed using the Visium Mouse Transcriptome Probe Set v2.0 (mm10-2020-A), supplemented with custom probes targeting RCAS-derived transgenes (one probe against human PDGFB and two probes against RFP). After hybridization, the labeled tissue was transferred to Visium HD slides using the Visium CytAssist instrument at the Center for Translational Genomics, Lund University.

#### Visium v1 10x Spatial Transcriptomics

For 10x Visium v1 spatial transcriptomics analysis, 2 primary and 2 recurrent snap-frozen tumors were processed. RNA Integrity Number (RIN)-values were >9 for all tumors. Tumor brain tissue had previously been embedded in OCT and snap-frozen in ice-cold isopentane and kept at -80°C until further processed. Performance and handling of slides for RNA extraction and library preparation were performed as recommended by manufacturer (Visium Spatial Gene Expression Reagent Kits, CG000239, Rev G, 10x Genomics). In brief, tissue was sectioned into 10 µm slices and placed on the pre-cooled 10x Genomics Visium Slide. The slides were fixed with ice-cold methanol, stained for Hematoxylin and Eosin (H&E). Images were acquired on Leica Dmi8 Microscope including the fiducial frame for further spot alignment. Tissue permeabilization was performed at 21 minutes (based on previous results from optimization slide of same tissue type). Second strand, cDNA amplification & Quality Control was performed according to the manufacturer’s user guide. Libraries were prepared and sequenced using NovaSeq 6000 SP (100 cycles) kit v1.5 at the Center for Translational Genomics, Lund University.

#### Spatial Data Processing

Raw sequencing data from all spatial transcriptomics experiments were processed using the 10x Genomics spaceranger software with platform-specific reference configurations. For Visium v1 datasets, spaceranger (v2.0.1) was used to align reads to a custom mouse reference genome generated using Cell Ranger mkref, based on the mm10-2020-A reference (refdata-gex-mm10-2020-A) and extended with Tva, human PDGFB and RFP genes. For Visium HD datasets, spaceranger (v3.1.3) was used with the Visium Mouse Transcriptome Probe Set v2.0 (mm10-2020-A) supplemented with custom probes targeting RCAS-derived transgenes (human PDGFB and RFP) as reference. Resulting spatial gene expression matrices and associated imaging data were imported into R and analyzed using Seurat.

For Visium HD, Spatial gene expression data were generated at 16 µm bin resolution. Module scores for curated gene signatures, including GBM and astrocyte subtype programs, were computed using Seurat’s AddModuleScore and visualized as spatial expression maps. For integrative analysis with the matched single-cell Flex dataset, Visium HD data were additionally binned at 8 µm resolution and subjected to label transfer using Seurat’s FindTransferAnchors and TransferData functions, yielding per-bin prediction scores for reference-derived cell state labels.

For Visium v1, two primary and two post-radiotherapy recurrent tumors were analyzed using the same curated gene signature collection. Spatial assays were log-normalized, variable features identified, and data scaled in Seurat. Module scores were computed using AddModuleScore. For each sample, Spearman correlation matrices were calculated across gene signatures. Using Neopl-ACR and Astrocyte_R as target signatures, correlations with all other signatures were summarized across samples by mean, minimum, and maximum values. These relationships were visualized as ranked bar plots with whiskers to summarize consistency and variability between primary and recurrent tumors.

### Visium v1 External human cohort analysis

Twenty Visium v1 GBM samples from the Heiland *et al*. dataset were processed by loading the filtered_feature_bc_matrix.h5 files, normalizing with NormalizeData, identifying variable features, scaling, and scoring humanized signatures with AddModuleScore. A Spearman correlation matrix was computed per sample. Samples were grouped by biological category based on sample IDs (tumor T/TC/TI, non-tumor cortex C, IDH-mutant gliomas). For each group, correlations of all signatures with Neopl-ACR or Astrocyte_R module scores were summarized by mean, minimum, and maximum values and plotted as ranked barplots to compare associations across biological states.

#### Software

All analyses were performed in R version 4.3.0 using Seurat v5.1, SeuratExtend, SCpubr, scCustomize, Matrix, ggplot2, ComplexHeatmap, AUCell, CellChat v1.6.1, LIANA v1.2.0, and associated tidyverse packages. The complete Seurat object (raw and processed) and sample-level metadata will be made available through public repositories as indicated in the Data Availability section.

## Data and code availability

Raw sequencing data and primary processing outputs (Cell Ranger and Space Ranger) have been deposited in ArrayExpress under accessions E-MTAB-16420, and E-MTAB-16421. Analysis-ready Seurat objects used for downstream analyses and figure generation are available via Zenodo (https://doi.org/10.5281/zenodo.17922854). All analysis code is available at GitHub (https://github.com/PietrasLab/gbm-rcas-2025-lindgren-rosberg/).

