## Supplementary Figures for "Coevolution of neoplastic and non-neoplastic reactive astrocyte states converges on mesenchymal-like and injury-response programs during murine glioblastoma progression and post-radiotherapy recurrence"

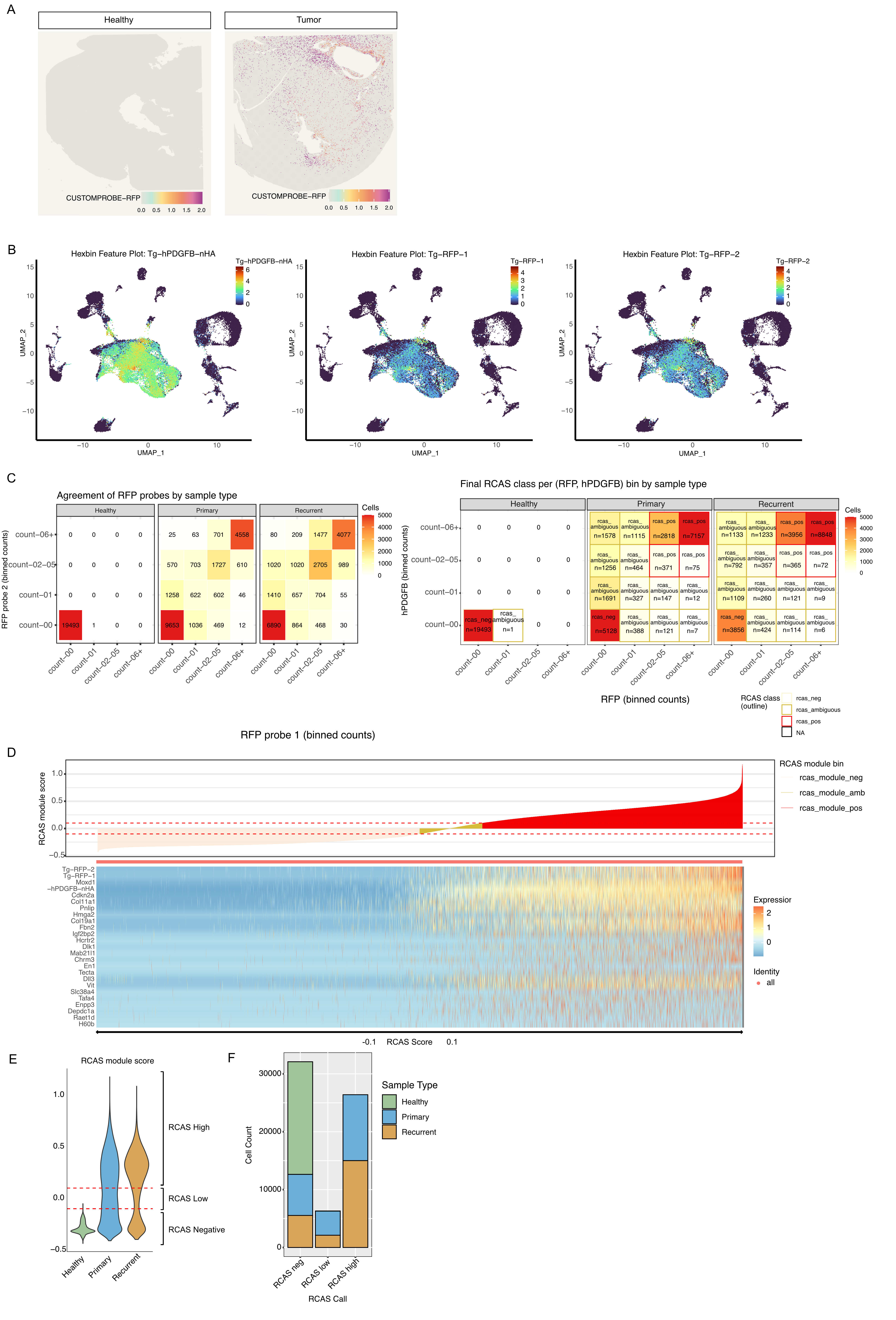

**Supplementary figure 1.** RCAS transgene–based identification of neoplastic cells by expression and module score

**(A)** Spatial transcriptomics feature plots (Visium v1) showing expression of the RCAS-delivered transgene RFP in healthy brain and tumor sections, demonstrating tumor-restricted detection of the viral transcript.

**(B)** UMAP feature plots from the scRNA-seq dataset (10x Genomics Flex; n = 64,804 cells) showing per-cell expression of the RCAS transgenes used in this study: Tg-hPDGFB-nHA and the two probes targeting RFP (Tg-RFP-1 and Tg-RFP-2). Transgene expression is restricted to tumor-derived cells and forms a coherent “RCAS+” compartment in the embedding. **C to F:** RCAS-based cell classification schema: RCAS transgene expression → joint hPDGFB/RFP expression bins → differential expression analysis → RCAS gene module score → final RCAS-based cell classification. **(C)** Intermediate transgene binning used to derive the RCAS signature. , agreement between the two RFP probe features across sample types (healthy, primary, recurrent), shown as binned transgene counts per cell. Right, joint expression bins derived from combined hPDGFB and RFP expression, integrating transgene detection across probes. These joint RCAS expression bins were used as the grouping variable for downstream differential expression analysis (Seurat FindAllMarkers) to identify genes enriched in RCAS-positive versus RCAS-negative cells. **(D)** Construction of the RCAS module score. Cells are ordered by the resulting RCAS module score (top; dashed lines indicate thresholds used to define RCAS^neg^, RCAS^low^, and RCAS^high^), together with a heatmap of the RCAS gene set derived from the joint-bin differential expression analysis (bottom). This module score captures coordinated transcriptional programs associated with RCAS-driven neoplastic cells and mitigates sparsity in individual transgene detection. **(E)** Distribution of RCAS module scores across biological groups (healthy, primary, recurrent), demonstrating tumor-specific enrichment of RCAS-low and RCAS-high cells. **(F)** Final RCAS classification across sample types. Cells were assigned as RCAS^high^, RCAS^low^, or RCAS^neg^ based primarily on the RCAS module score. As a final correction step, cells exhibiting high transgene expression but low RCAS module score were reassigned as RCAS-positive (286 of 64,804 cells).

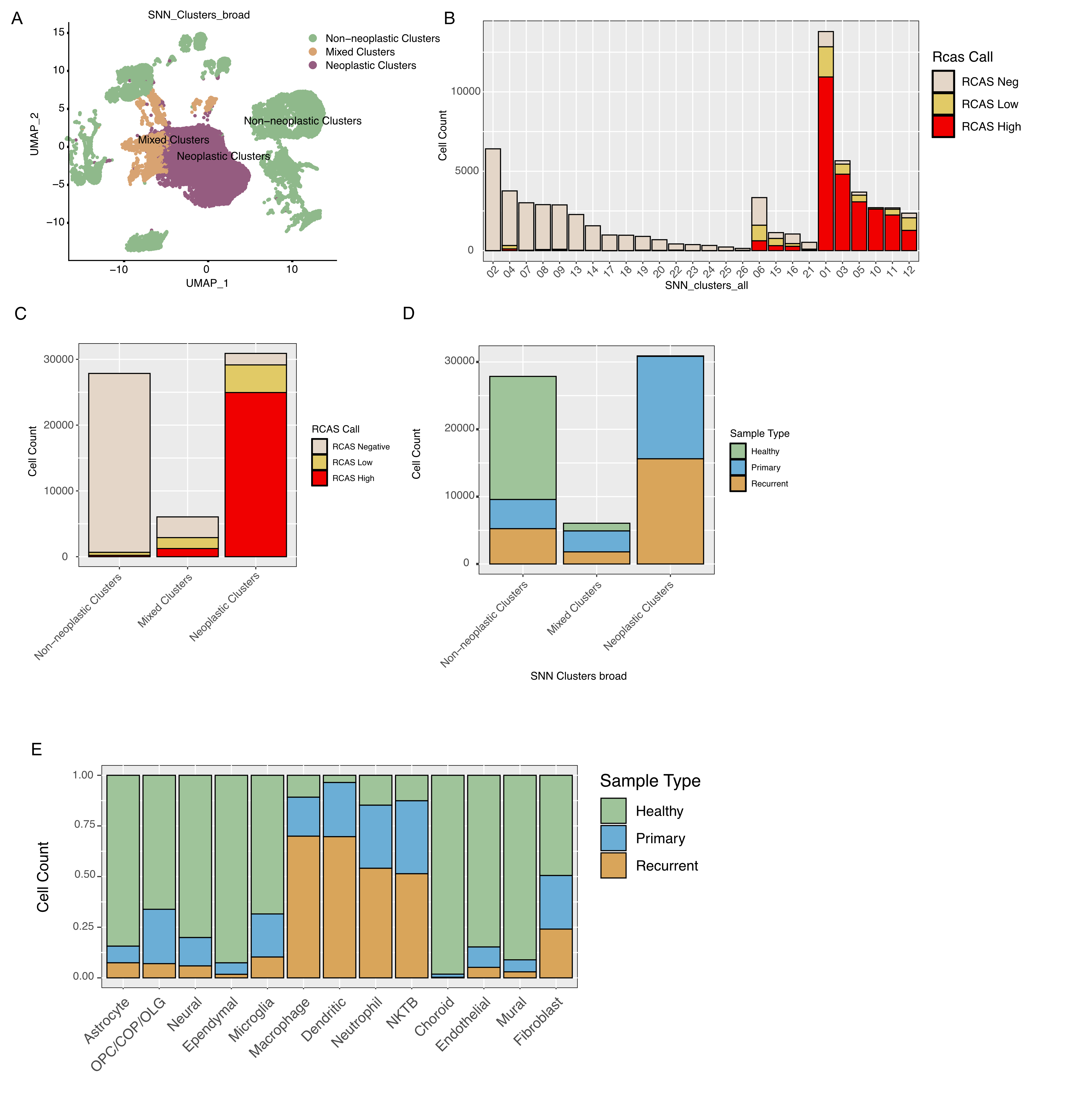

**(A)** UMAP projection of all 64,804 single cells showing the same 26 SNN clusters as in Figure 2A, here colored according to a broad RCAS-based grouping of those clusters into non-neoplastic, mixed, and neoplastic compartments. This grouping reflects RCAS transgene expression patterns and does not represent an additional clustering step. **(B)** Distribution of RCAS expression status (RCAS^neg^, RCAS^low^, RCAS^high^) across the 26 individual SNN clusters. Clusters are ordered according to their assignment to non-neoplastic, mixed, or neoplastic groups, illustrating how cluster-level RCAS composition underlies the broad classification shown in (A). **(C)** Aggregated RCAS expression status within each broad SNN group. Non-neoplastic clusters are dominated by RCAS^neg^ cells, neoplastic clusters consist almost exclusively of RCAS^low/high^ cells, and mixed clusters contain both populations. **(D)** Distribution of sample types (healthy, primary tumor, recurrent tumor) across the three broad SNN groups. Neoplastic clusters are composed exclusively of tumor-derived cells, whereas mixed clusters contain contributions from all sample types. **(E)** Relative abundance of major non-neoplastic cell types across healthy tissue, primary tumors, and recurrent tumors.

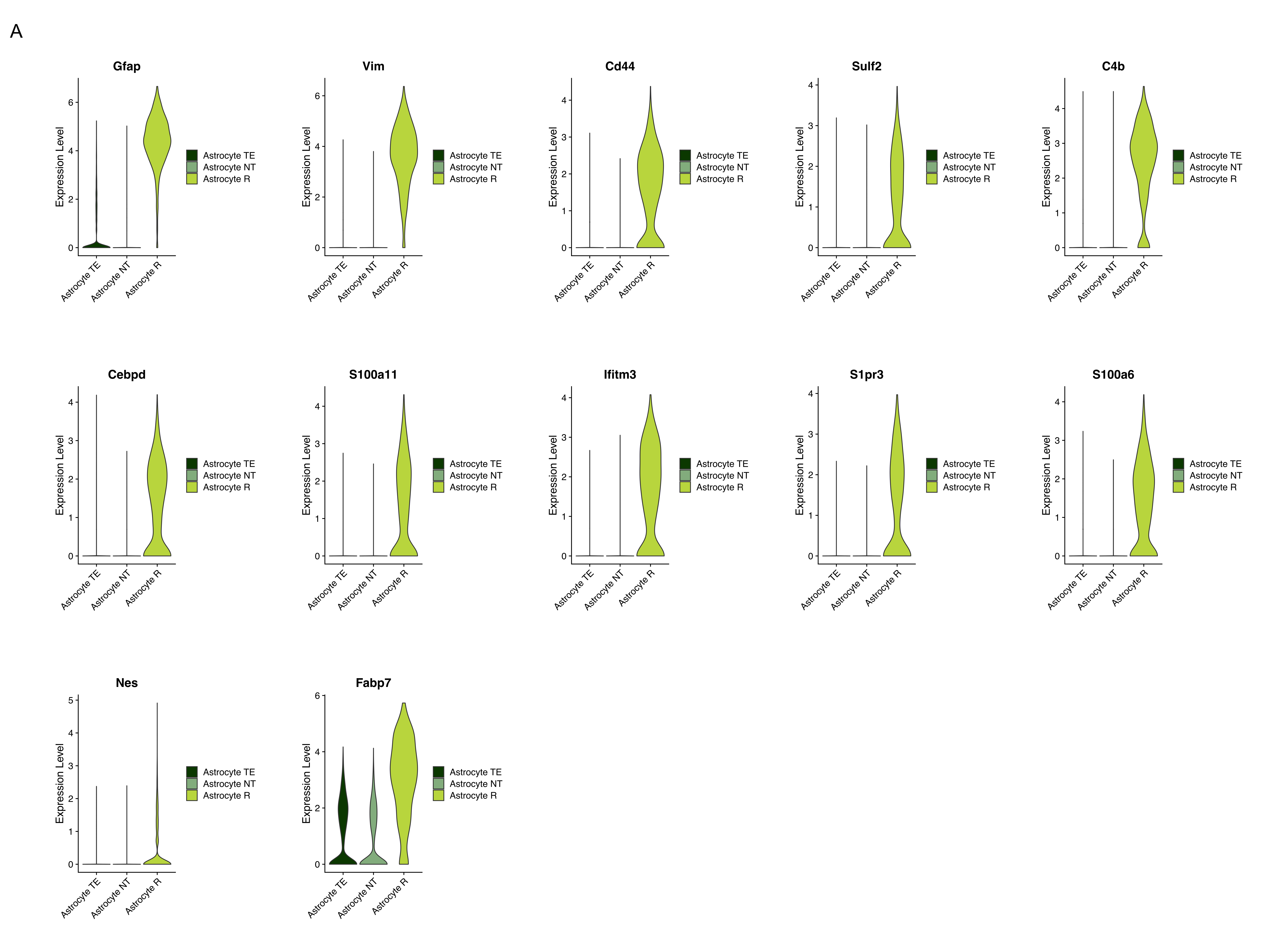

**Supplementary figure 3.**

**(A)** Gene expression level of astrocyte reactivity and inflammation markers (*Gfap*, *Vim*, *Cd44*, *Sulf2*, *C4b*, *Cebpd*, *S100a11*, *Ifitm3* and *S1pr3*) and astrocytic progenitor cells (*Gfap*, *Vim*, *Cd44*, *S100a6*, *Nes*, *Fabp7*).

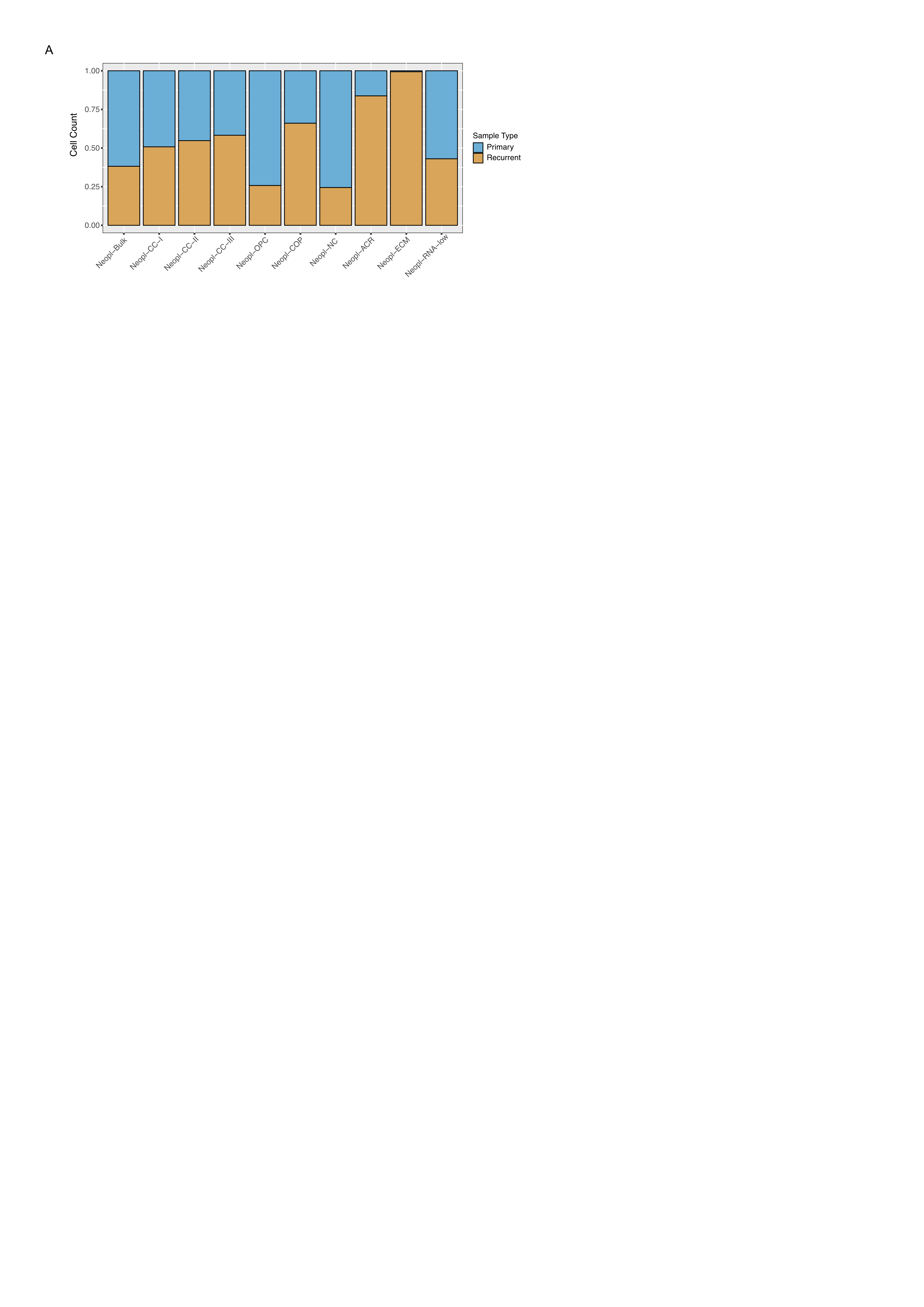

**Supplementary figure 4.**

**(A)** Relative abundance of major non-neoplastic cell types found in primary and recurrent sample types. Recurrent tumors exhibit expansion of Neopl-ECM and Neopl-ACR, whereas Neopl-OPC and Neopl-NC decrease.

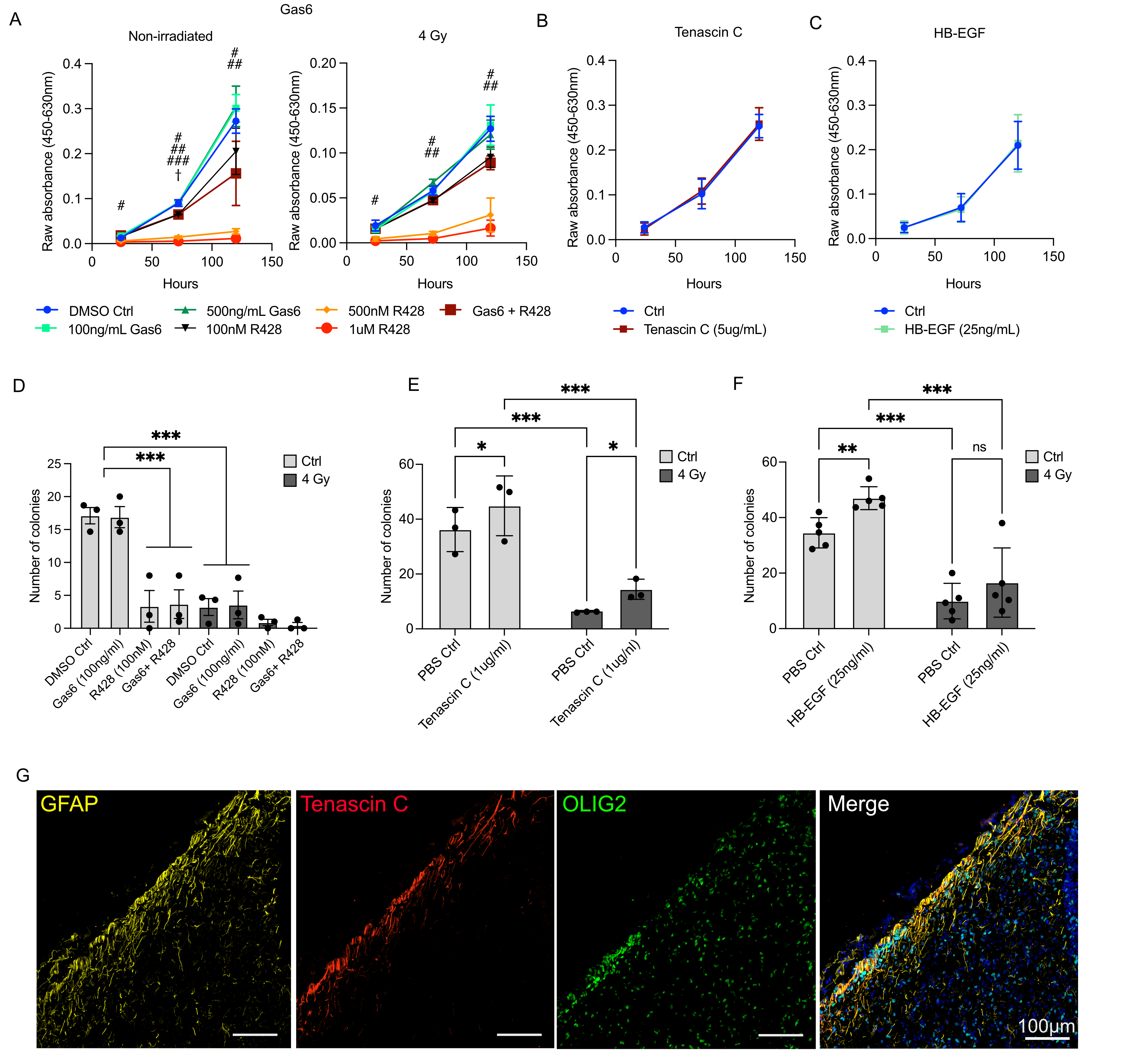

**Supplementary figure 5.**

(A) Proliferation of PDGFB-induced glioma primary cultures (PIGPC) in presence of recombinant Gas6 or Axl inhibitor R428 with or without 4 Gy radiotherapy. All groups were compared to DMSO Ctrl within each timepoint and statistical analysis was performed with One-Way ANOVA with Dunnett’s multiple comparisons test where ‘#’ = DMSO Ctrl versus 1uM R428, ‘##’ = DMSO Ctrl versus 500nM R428, ‘###’ = DMSO Ctrl versus 100nM R428, ‘†’ = DMSO Ctrl vs. 100nM R428 + 100ng/ml Gas6. Comparisons with P-values < 0.05 are shown in the graph. (B-C) PIGPC cell proliferation in presence of recombinant Tenascin C or HB-EGF as measured by WST-1 assay. (D-F) Quantification of clonal survival of PIGPC in presence of recombinant Gas6 and Axl inhibitor R428, recombinant Tenascin C or HB-EGF with or without 4 Gy radiotherapy, statistical analysis was performed with Two-way ANOVA with Sidak’s multiple comparison for (D) and Uncorrected Fisher’s LSD test for (E-F). (G) Gene expression of Gfap, Gas6, Tnc and Hbegf mapped on spatially resolved transcriptomics from healthy (G) and tumor (H) tissue generated with RCAS-tv-a. (I) Representative image showing immunofluorescent staining of GFAP (yellow), Tenascin C (red) and OLIG2 (green) on a murine glioma. Statistical significance based on *P < 0.05, **P < 0.01 and ***P< 0.001.

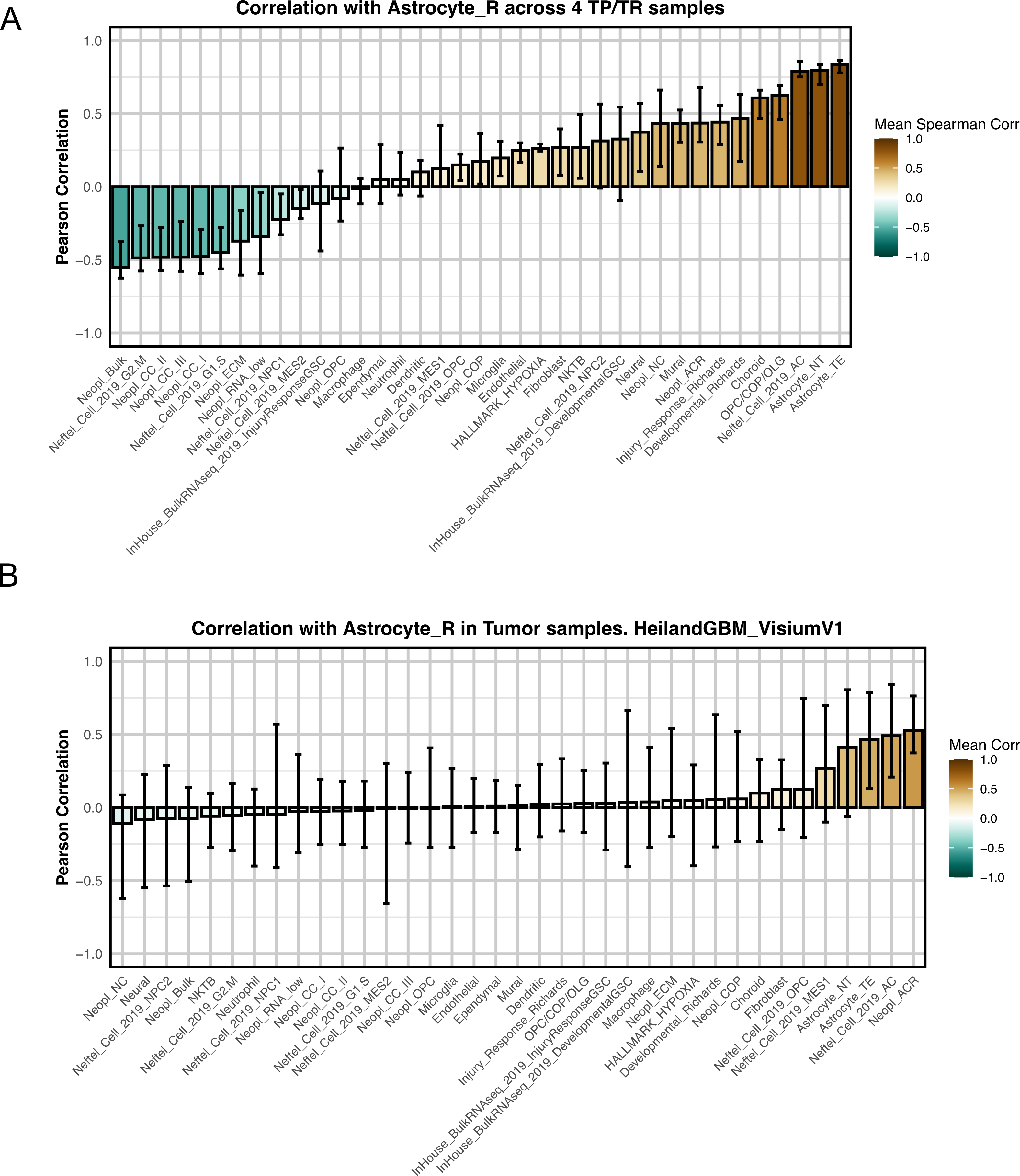

**Supplementary figure 6.**

**(A)** Ranked mean Spearman correlations between the Astrocyte_R module score and all other signatures across four in-house Visium v1 samples. Bars show the mean correlation across the four samples; error bars indicate the minimum and maximum correlation observed. **(B)** As in (A), but for human GBM Visium v1 data from Heiland *et al*., restricted to tumor samples. Astrocyte R correlations are consistently highest with Astrocyte NT, and TE, Neopl_ACR and Neftel_Cell_2019_AC.
